# Coordinate control of the RNA polymerase II transcription cycle by CDK9-dependent, tripartite phosphorylation of SPT5

**DOI:** 10.1101/2024.07.25.605161

**Authors:** Rui Sun, Robert P. Fisher

## Abstract

The RNA polymerase II (RNAPII) transcription cycle is regulated throughout its duration by reversible protein phosphorylation. The elongation factor SPT5 contains two regions targeted by cyclin-dependent kinase 9 (CDK9) and previously implicated in promoter-proximal pausing and termination: the linker between KOWx-4 and KOW5 domains and carboxy-terminal repeat (CTR) 1, respectively. Here we show that phosphorylations in the KOWx-4/5 linker, CTR1 and a third region, CTR2, coordinately control pause release, elongation speed and RNA processing. Pausing was increased by mutations preventing CTR1 or CTR2 phosphorylation, but attenuated when both CTRs were mutated. Whereas mutating CTR1 alone slowed elongation and repressed nascent transcription, simultaneous mutation of CTR2 partially reversed both effects. Nevertheless, mutating both CTRs led to aberrant splicing, dysregulated termination and diminished steady-state mRNA levels, and impaired cell proliferation more severely than did either single-CTR mutation. Therefore, tripartite SPT5 phosphorylation times pause release and regulates RNAPII elongation rates positively and negatively to ensure productive transcription and cell viability.

## INTRODUCTION

The transcription cycle of RNA polymerase II (RNAPII) comprises initiation, elongation, termination, and the recycling of enzymatic machinery to permit new rounds of initiation ^1–3^. Transitions between phases are regulated by phosphorylation of RNAPII and its accessory factors, which can be rate-limiting for expression of certain genes ^4,5^. More generally, reversible phosphorylation imposes quality control—coupling synthesis and co-transcriptional processing of RNA ^6,7^, coordinating progress of elongating RNAPII with chromatin modification ^2,8^, and ensuring efficient termination to allow RNAPII recycling and avoid transcriptional interference between neighboring genes ^9,10^. This coordination is critical for normal development and frequently disrupted in cancer cells ^11,12^, in which transcriptional derangement can be a potential vulnerability; the kinases and phosphatases that govern the RNAPII transcription cycle have emerged as promising targets for therapeutic intervention ^12–15^.

The RNAPII transcription complex undergoes a structural transformation shortly after initiation, with the exchange of initiation factors for elongation factors—the DRB sensitivity-inducing factor (DSIF) composed of SPT4 and SPT5, and the multi-subunit negative elongation factor (NELF) ^1^. In human cells, this exchange depends on the catalytic activity of cyclin-dependent kinase (CDK) 7 ^16–20^—part of the initiation factor TFIIH ^21,22^—and, at many genes, results in a stable, promoter-proximal pause in elongation within ∼100 nucleotides (nt) of the transcription start site (TSS) ^5^. Pausing regulates expression of genes involved in cell division, signal-response pathways, and development ^4,5,23,24^. Pause release depends on CDK9, catalytic subunit of positive transcription elongation factor b (P-TEFb), which phosphorylates multiple components of the paused elongation complex, including RNAPII, SPT5 and NELF, to trigger NELF eviction and conversion of DSIF into a positive elongation factor ^5,25^.

Besides RNA polymerase itself, DSIF is the only component of the transcription machinery conserved in all domains of life, with orthologs in prokaryotes and archaea ^26^. In the fission yeast *Schizosaccharomyces pombe*, Spt5 is globally required for transcript elongation and viability ^27^, and genetic loss of Spt4, while not lethal, attenuated promoter-proximal pausing ^28^. Recent studies in human cells uncovered a role for DSIF in stabilizing RNAPII; acute depletion of SPT5 led to ubiquitin-dependent degradation of chromatin-associated RNAPII ^29,30^.

SPT5 is a modular protein with a carboxy-terminal region harboring repetitive sequences phosphorylated by CDK9 ^31–33^. In *S. pombe*, phosphorylation in the Spt5 carboxy-terminal domain (CTD) was enriched during elongation and removed downstream of the cleavage and polyadenylation signal (CPS), where RNAPII pauses again in preparation for termination ^34^. In both yeast and human cells, CDK9 and phosphoprotein phosphatase 1 (PP1) work in opposition to control SPT5 carboxy-terminal phosphorylation, RNAPII elongation speed and termination ^34–38^.

Metazoan SPT5 is also phosphorylated by CDK9 on a flexible linker between Kyrpides-Ouzounis-Woese (KOW) motifs x-4 and 5 ^15,25,39^; this region binds nascent RNA in RNAPII elongation complexes ^40,41^ and is required for promoter-proximal pausing *in vitro* ^42,43^. The phosphorylated KOWx-4/5 linker is a substrate for two related protein phosphatases—PP4 and the PP2A module of the Integrator complex—that are both implicated in regulating RNAPII elongation in opposition to CDK9 ^14,38,44–46^. Genetic dissections of human SPT5 indicated a role for KOWx-4/5 linker phosphorylation in release from the promoter-proximal pause ^30,47^, and for carboxy-terminal repeat region (CTR) 1 phosphorylation in termination; introduction of a mutant variant with all seven Thr residues in CTR1 changed to Ala shortened the distance transcribed downstream of the CPS before termination ^30^.

Vertebrate SPT5 contains a second region nearer the carboxyl terminus, CTR2 ^31^, with 22 sites matching the minimal consensus motif for CDK phosphorylation, S/T-P ^39,48–50^. In vitro, CDK9 phosphorylated CTR2 less efficiently than it did CTR1 ^51^, and previous analyses suggested CTR2 was dispensable, or redundant with CTR1, for SPT5 function ^52^. Here we show that CTR2 is phosphorylated in human cells, and becomes more heavily modified when CTR1 phosphorylation is blocked, suggesting communication, or competition for CDK9 activity, between the CTRs. Whereas mutation of either CTR increased promoter-proximal RNAPII occupancy, mutating both attenuated pausing, which was restored by additional mutation of the KOWx-4/5 linker. However, CTR1 and CTR2 are not mechanistically redundant; preventing CTR1 phosphorylation slowed elongation and repressed nascent transcription, while simultaneous mutation of CTR2 partially restored elongation velocity and increased transcription of a subset of mostly pause-regulated genes. Whereas CTR1 and CTR2 had opposing effects on elongation rate, they had reinforcing effects on gene expression and viability. The double-CTR mutation had pervasive effects on mRNA maturation, with altered splicing and aberrant termination affecting different gene subsets, and caused greater repression of mRNA levels and more severe growth defects than did either single-CTR mutation. We propose that phosphorylations in CTR1 and CTR2 of SPT5 perform a timing function in pause release, and act on elongating RNAPII as accelerator and brake, respectively, to coordinate RNA processing and termination and so ensure productive transcription.

## RESULTS

### A chemical genetic system for functional dissection of SPT5

To uncover the functions of SPT5 phosphorylation in human cells, we sought to deplete the endogenous protein and replace it with variants mutated at known or suspected sites of phosphorylation. Our initial attempts using RNA interference (RNAi) achieved ∼70% reduction in SPT5 levels in whole-cell extracts of human HCT116 colon cancer cells, but little or no loss of chromatin-bound SPT5 or RNAPII, or of phosphorylation in CTR1 or at Ser666 in the KOWx-4/5 linker (**Figures S1A** and **S1B**), suggesting that these cells maintain an ample reservoir of free SPT5 and robust mechanisms to preserve SPT5 function in transcription.

To circumvent these homeostatic pathways, we turned to an inducible-degron (dTAG) strategy ^53^. We introduced, by CRISPR/Cas9-mediated genome editing, a cassette encoding a hemagglutinin (HA) epitope and the FKBP12^F36V^ degron into both copies of the *SUPT5H* gene encoding SPT5 (**Figure S1C**). Accurate, homozygous integration was confirmed by PCR and sequencing (**Figures S1D** and **S1E**). Immunoblot analysis of wild-type and *dTAG-SUPT5H* cell extracts revealed a shift of SPT5 to slower mobility in the latter, consistent with the larger expected size of the tagged protein, and depletion of dTAG-SPT5 but not wild-type SPT5 upon addition of the small-molecule inducer dTAGv-1 (**Figure 1A**). The degradation of dTAG-SPT5 was both time- and dose-dependent, with SPT5 becoming undetectable within 1 hr of treatment with 125 nM dTAGv-1 (**Figures 1B** and **S1F**). Critically, degradation of soluble and chromatin-associated dTAG-SPT5 was similarly rapid and efficient (**Figure 1B**). Consistent with loss of SPT5 function in transcription, chromatin immunoprecipitation and sequencing (ChIP-seq) analysis revealed globally diminished RNAPII occupancy, most markedly in promoter-proximal regions, upon depletion of dTAG-SPT5 (**Figure 1C**). Degradation of dTAG-SPT5 led to concomitant depletion of RPB1, the largest subunit of RNAPII (**Figure 1D**), presumably reflecting the previously reported role of SPT5 in protecting RNAPII from ubiquitin-dependent degradation ^29,30^.

**Figure 1.**
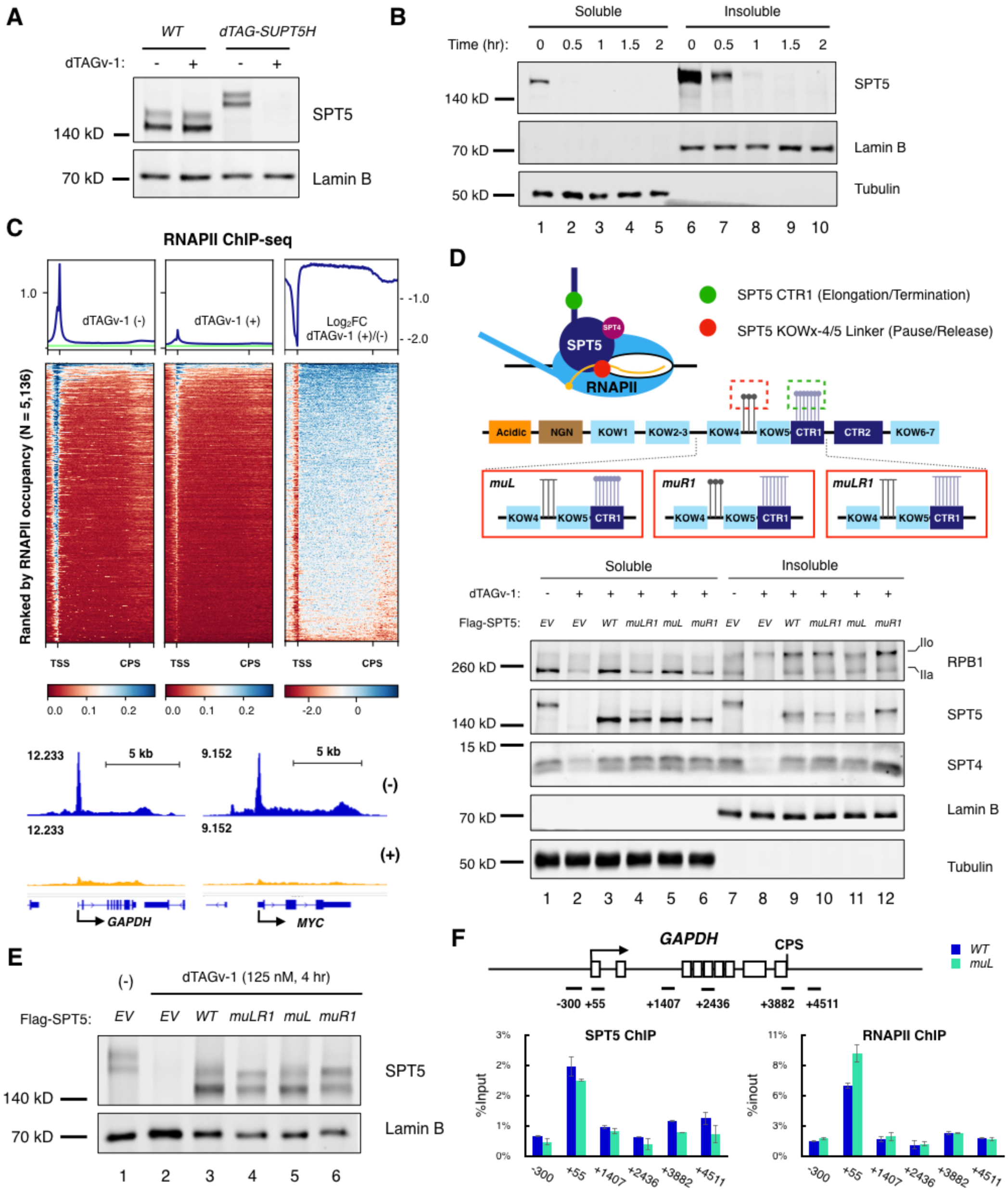
A chemical genetic system for functional dissection of SPT5. **A.** Wild-type (*WT*) or *dTAG-SUPT5H* cells, as indicated, were treated with with DMSO (-) or 0.5 µM dTAGv-1 (+) for 4 hr and immunoblotted with indicated antibodies. **B.** Soluble and insoluble (chromatin-associated) fractions of *dTAG-SUPT5H* cells were analyzed by immunoblotting with indicated antibodies after treatment with 125 nM dTAGv-1 for indicated times. **C.** ChIP-seq analysis in *dTAG-SUPT5H* cells treated with (+) or without (-) dTAGv-1 (125 nM, 4 hr). Spike-in normalized metagene plots (top), heatmaps (middle), and browser tracks (bottom) of RNAPII (RPB1) were merged from two biological replicates. Green line: intergenic signal. **D.** Replacement of endogenous dTAG-SPT5 with Flag-SPT5 variants. Top: Cartoon representation of different phosphorylated regions of SPT5 in relation to RNAPII and nascent RNA. Middle: Different Flag-SPT5 mutant variants ectopically expressed in *dTAG-SUPT5H* cells (*muL, muR1, and muLR1*). Sticks with blunt endpoints indicate Ala substitutions, while round endpoints indicate Ser or Thr residues. Bottom: *dTAG-SUPT5H* cells expressing indicated Flag-SPT5 variants were treated with DMSO or 125 nM dTAG-v1, as indicated, for 4 hr, fractionated and analyzed by immunoblotting with indicated antibodies. *EV*: empty vector control. **E.** Immunoblot analysis of Flag-SPT5 variants expressed in *dTAG-SUPT5H* cells in formaldehyde cross-linked chromatin fractions. Fractions from indicated conditions were analyzed after cross-link reversal. **F.** ChIP-qPCR of SPT5 and RNAPII (RPB1) at *GAPDH* gene in *dTAG-SUPT5H* cells expressing *WT* or *muL* Flag-SPT5 variant after dTAGv-1 treatment (125 nM, 4 hr). Data are from two independent experiments (n = 2); error bars indicate mean ± SD.

Next, to assess contributions of phosphorylation to SPT5 function, we stably expressed Flag-tagged SPT5 variants in *dTAG-SUPT5H* cells. The initial set (**Figure 1D**) included wild-type SPT5 (*WT*) and variants with Ala substitutions of 1) Ser666, Ser671 and Ser686, three sites phosphorylated by CDK9 ^25^ within the KOWx-4/5 linker (*muL*); 2) Thr residues at the fourth position of all seven CTR1 repeats (consensus: G_1_-S_2_-Q/R_3_-T_4_-P_5_-X_6_-Y_7_ where X is any residue ^31^) (*muR1*); and 3) all ten potential phosphorylation sites in both the KOWx-4/5 linker and CTR1 (*muLR1*). All variants were expressed at levels similar to that of endogenous dTAG-SPT5, and prevented depletion of RPB1 and the SPT5 binding partner SPT4 (**Figure 1D**). Mutation of the KOWx-4/5 linker, however, diminished recovery of SPT5 in a bulk chromatin fraction (**Figure 1D**, compare lanes 10 and 11 to lanes 9 and 12) and reduced Flag-SPT5 signals detected by cleavage under targets and release using nuclease (CUT&RUN) (**Figure S1G**)—effects reminiscent of a previous study in which deletion of KOW4 and KOW5 reduced SPT5 association with *Drosophila* polytene chromosomes ^42^. In formaldehyde cross-linked chromatin, however, recovery of *muL* and *muLR1* was close to that of *WT* and *muR1* variants, respectively (**Figure 1E**), and ChIP-qPCR analysis revealed similar occupancy by *WT* and *muL* variants (**Figure 1F**). Taken together, the results suggest that an intact KOWx-4/5 linker contributes to stability, but not establishment, of SPT5 association with chromatin.

### SPT5 CTR2 is phosphorylated in human cells

We next probed immunoblots of the different variants with previously characterized phospho-specific antibodies ^38,39^. Reactivity with an antibody specific for phosphorylated CTR1 (pCTR1) was nearly abolished in the CTR1-mutant variants but unaffected, relative to signals with a pan-specific SPT5 antibody, by mutations in the KOWx-4/5 linker (**Figure 2A**). Therefore, CTR1 phosphorylation appears to be independent of KOWx-4/5 linker phosphorylation, consistent with ChIP-seq analysis, in which pCTR1 accumulated upstream of phosphorylated Ser666 (pSer666) ^38^. On the other hand, an antibody that recognizes pSer666 had increased reactivity with the *muR1* variant, but this was apparently not due to increased KOWx-4/5 linker phosphorylation; the *muLR1* variant also produced a stronger signal than *muL* (**Figure 2A**). We suspected that anti-pSer666 might recognize phosphorylated CTR2 (pCTR2), which contains 22 S/T-P sites, with several near matches to the immunogenic peptide, VGGFAPM[pS]PRISSP, used to raise the antibody ^39^ (**Figures 2A** and **S2A**). We therefore expressed a second set of Flag-SPT5 variants in *dTAG-SUPT5H* cells with an internal deletion of CTR2 (residues 841-968). With CTR2 removed, anti-pSer666 signals became strictly dependent on an intact KOWx-4/5 linker and unaffected by CTR1 mutations (**Figure 2B**), suggesting that the increased phosphorylation detected by anti-pSer666 when CTR1 phosphorylation was blocked occurred in CTR2.

**Figure 2.**
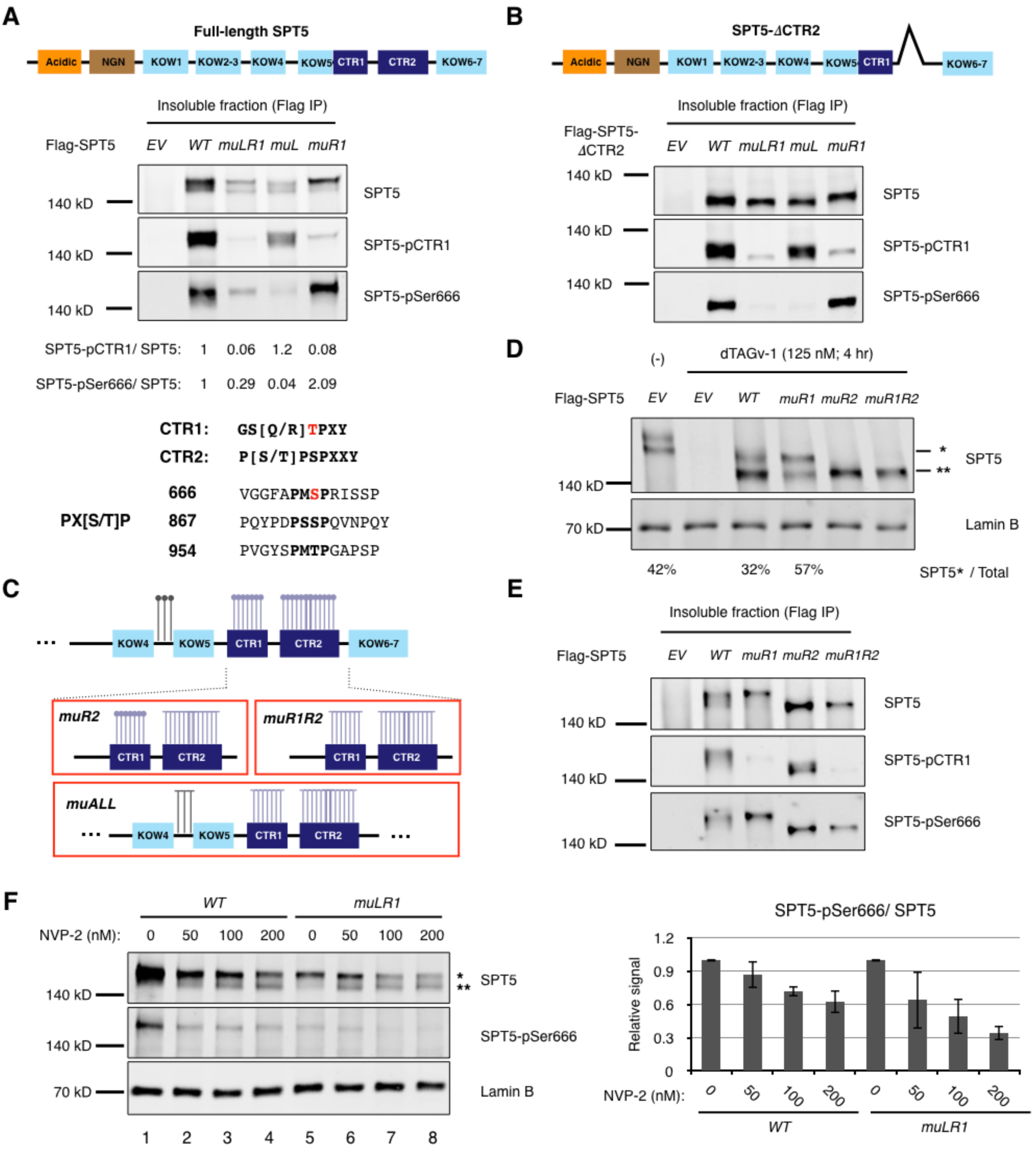
SPT5 CTR2 is phosphorylated in human cells. **A.** Top: Schematic of full-length SPT5 with conserved motifs indicated. Middle: Immunoblot analysis of indicated, full-length Flag-SPT5 variants expressed in *dTAG-SUPT5H* cells treated with DMSO (-) or dTAGv-1 (125 nM, 4 hr), as indicated, immunoprecipitated with anti-Flag antibodies and probed with indicated pan- and phospho-specific SPT5 antibodies. Numbers under lanes indicate ratio of phospho-SPT5 to total SPT5 signals in variants, relative to ratio in wild-type (defined as 1). Bottom: Consensus sequences of CTR1 and CTR2 repeats, and sequence surrounding Ser666 compared to two CTR2 repeats. **B.** Top: Schematic of SPT5 with internal deletion of CTR2. Bottom: Immunoblot analysis of indicated Flag-SPT5 variants with internal deletion of CTR2, expressed in *dTAG-SUPT5H* cells treated with DMSO (-) or dTAGv-1 (125 nM, 4 hr), as indicated, immunoprecipitated with anti-Flag antibodies, and probed with indicated pan- and phospho-specific SPT5 antibodies. **C.** Schematic of SPT5 CTR2-mutant variants (*muR2, muR1R2, and muALL)*. Sticks with blunt endpoints indicate Ala substitutions, while round endpoints indicate Ser or Thr residues. Note: number of potential phosphosites we mutated in CTR2 (22) is larger than the number depicted. **D.** Immunoblot of Flag-SPT5 variants expressed in *dTAG-SUPT5H* cells in formaldehyde cross-linked chromatin fractions, probed with anti-SPT5 antibody. Fractions were analyzed after cross-link reversal. Asterisks (* and **) indicate two electrophoretic isoforms of SPT5. Relative abundance of slower-migrating form (*/[* + **]) is expressed as a percentage under select lanes. **E.** Immunoblot of indicated, chromatin-associated Flag-SPT5 variants expressed in *dTAG-SUPT5H* cells treated with DMSO (-) or dTAGv-1 (125 nM, 4 hr), as indicated, immunoprecipitated with anti-Flag antibodies and probed with indicated pan- and phospho-specific SPT5 antibodies. **F.** Immunoblot of indicated, chromatin-associated Flag-SPT5 variants, expressed in *dTAG-SUPT5H* cells treated with 125 nM dTAGv-1 for 4 hr and indicated doses of NVP-2 for 1 hr, probed with indicated antibodies. The ratios of pSer666: total SPT5 signals are plotted at right. Data are from three independent experiments (n = 3); error bars indicate mean ± SD.

To investigate CTR2 phosphorylation further, we expressed another set of Flag-SPT5 mutant variants (**Figure 2C**) with 1) all 22 S/T-P sites in CTR2 changed to A-P (*muR2*); 2) all such sites in both CTRs mutated (*muR1R2*); and 3) the KOWx-4/5 linker, CTR1 and CTR2 all mutated (*muALL*) (**Figures 2D** and **S2B**). Mutation of CTR2 altered SPT5 electrophoretic mobility; whereas wild-type and CTR1-mutant SPT5 each ran as a doublet, the *muR2* and *muR1R2* variants generated single bands at the faster-migrating position, which retained pSer666 and, in the case of *muR2*, pCTR1 (**Figure 2E**). Treatment with λ phosphatase collapsed SPT5 in immunoprecipitates of wild-type, *muR1* and *muLR1* variants into a single, fast-migrating band (**Figure S2C**), while mutation of CTR1 caused redistribution into the slower-migrating species (**Figure 2D**, compare *WT* to *muR1*), consistent with increased CTR2 phosphorylation. Taken together, the data suggest that CTR2 phosphorylation is both necessary and sufficient for the mobility shift, and that chromatin-associated SPT5 is normally phosphorylated in all three regions: the KOWx-4/5 linker, CTR1 and CTR2. Moreover, CTR2 phosphorylation, while underrepresented in previous phosphoproteomic studies (**Figure S2D**), occurs in ∼one-third of wild-type SPT5 (**Figure 1A, 1E** and **2D**), and is increased by mutations that prevent CTR1 phosphorylation, suggesting communication—or competition for a kinase—between the CTRs.

Phosphorylation of both CTR1 and Ser666 was catalyzed by CDK9 in whole-cell extracts ^39^ and sensitive to treatment with NVP-2, a selective inhibitor of CDK9 ^13,54^, in HCT116 cells ^38^. CDK9 also phosphorylated a substrate containing CTR2 fused to glutathione-*S*-transferase (GST) in vitro, albeit less efficiently than it did a GST-CTR1 fusion protein ^51^. To test whether phosphorylation of CTR2 in full-length SPT5 is CDK9-dependent, we treated cells expressing *WT* and *muLR1* variants with increasing concentrations of NVP-2, and assessed CTR2 phosphorylation by relative amounts of SPT5 electrophoretic isoforms in chromatin fractions, and by probing with the anti-pSer666 antibody, which reports exclusively on CTR2 phosphorylation when the KOWx-4/5 linker is mutated (**Figure 2F**, recovery of total SPT5 is decreased by the linker mutation as noted above). NVP-2 treatment caused a dose-dependent shift of both *WT* and *muLR1* variants to faster mobility. Moreover, the CTR2-dependent signals detected by anti-pSer666 with the *muLR1* variant were sensitive to NVP-2 at concentrations chosen to favor CDK9-selectivity ^13^. Therefore, phosphorylation of CTR2—like that of Ser666 and CTR1—is likely to depend at least in part on CDK9 in human cells.

### Reinforcing functions of CTR1 and CTR2 in gene expression and cell viability

To determine the impact of SPT5 phosphorylation-site mutations on cell proliferation and metabolic activity, we treated *dTAG-SUPT5H* cells with dTAGv-1 or vehicle and monitored cell numbers (**Figures 3A** and **S3A**), conversion of resazurin to resarufin (**Figure 3B**) and ATP levels (**Figure S3B**). Without ectopic expression of Flag-SPT5, cell proliferation and metabolism underwent rapid cessation, consistent with SPT5 being essential for viability ^55^. Expression of wild-type Flag-SPT5 restored normal proliferation with a doubling time < 24 hr. Mutation of CTR1 or CTR2 alone had moderate or negligible effects on this rescue, respectively, whereas their combined mutation severely impaired viability and metabolic activity (**Figures 3A, 3B** and **S3B**). Impaired proliferation was dependent on loss of endogenous dTAG-SPT5; without dTAGv-1 addition, growth rates of cells expressing different Flag-SPT5 variants (or no Flag-SPT5) were indistinguishable (**Figures 3A** and **S3A**).

**Figure 3.**
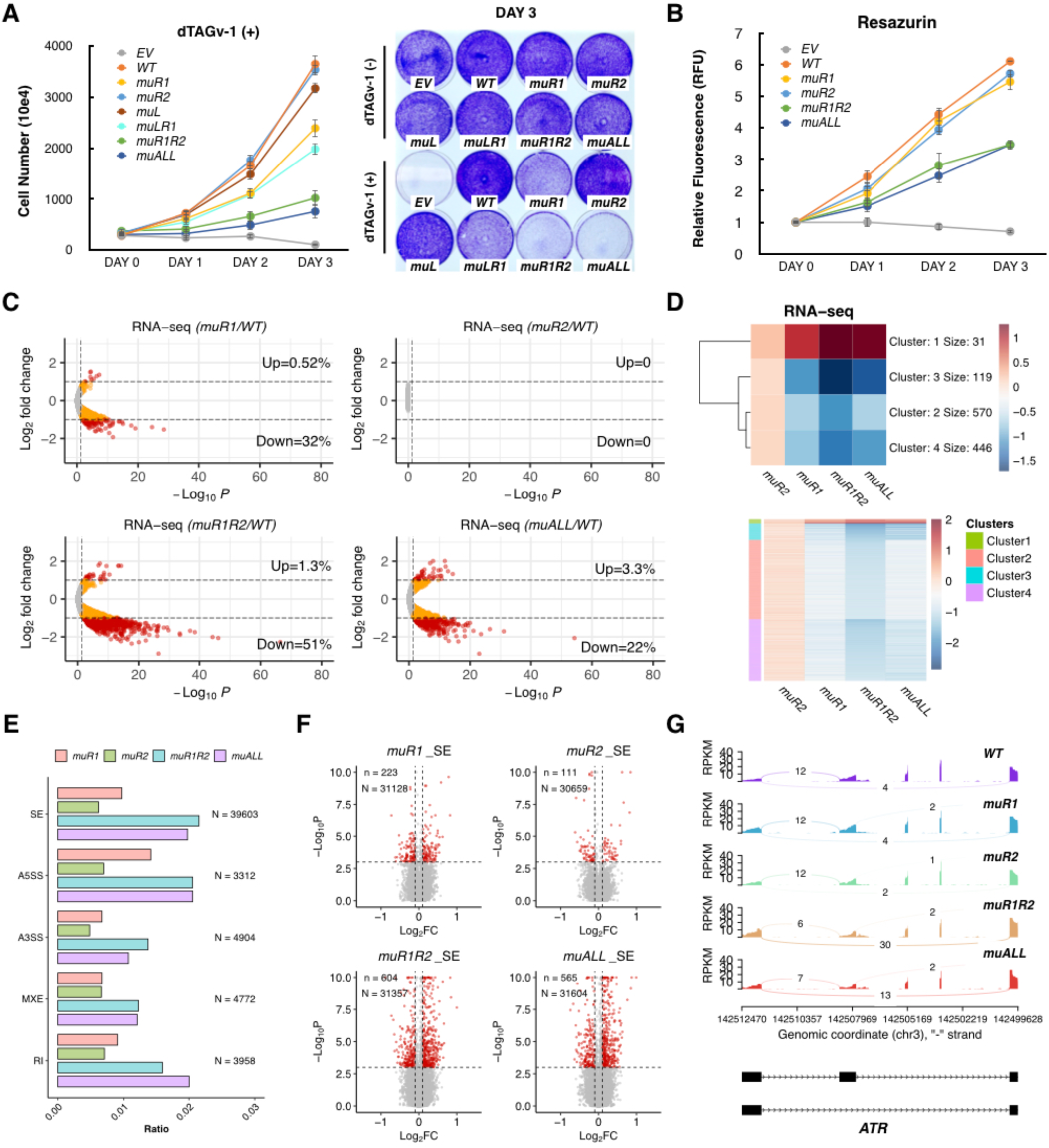
Reinforcing functions of CTR1 and CTR2 in gene expression and cell viability. **A.** Numbers of *dTAG-SUPT5H* cells expressing indicated Flag-SPT5 variants were counted on indicated days after treatment with 500 nM dTAGv-1 (left). Data are from three independent experiments (n = 3); error bars indicate mean ± SD. Representative crystal violet staining on day 3 is shown at right. **B.** Resazurin-to-Resarufin conversion by cells expressing indicated Flag-SPT5 variants after treatment with 500 nM dTAGv-1 for indicated times. Data are from four biological replicates (n = 4) for each time point; error bars indicate mean ± SD. **C.** Volcano plots of spike-in RNA-seq analysis in cells expressing indicated Flag-SPT5 variants after dTAGv-1 treatment (500 nM, 24 hr). Colored dots represent significantly changed genes in differential gene analysis (Wald test, adjusted p < 0.05, n = 10,383). Red dots indicate genes with fold-change > 2. Data are from two biological replicates (n = 2). **D.** Heatmap of RNA-seq data for genes most affected by double-CTR mutation (adjusted p < 0.05, fold change ≥ 2), aggregated into four clusters by k means clustering. **E.** Alternative splicing analysis. The ratio of significantly altered splicing events to total splicing events (N) is plotted on *x*-axis for Flag-SPT5 variants indicated on *y*-axis (likelihood-ratio test, FDR < 0.001, ΔIncLevel ≥ 0.1). Altered splicing events detected are: skipped exons (SE), alternative 5’ splices sites (A5SS), alternative 3’ splice sites (A3SS), mutually exclusive exons (MXE), and retained introns (RI). **F.** Volcano plots of exon-skipping (SE) in indicated mutants. Red dots represent significantly changed SE events (FDR < 0.001, ΔIncLevel ≥ 0.1, Counts ≥ 10). **G.** Sashimiplot of significantly changed SE events at the indicated gene locus.

We hypothesized that effects of *SUPT5H* mutations on cell proliferation would be reflected in steady-state mRNA levels, which we analyzed by RNA sequencing (RNA-seq) (**Figures 3C, 3D** and **S3C**). Mutating CTR1 alone decreased expression of ∼32% of genes (adjusted *p* < 0.05), of which 143 genes were repressed twofold or more, whereas CTR2 mutation alone globally but modestly increased mRNA levels, with no significantly repressed or induced genes (adjusted *p* < 0.05). When CTR1 and CTR2 mutations were combined, 51% of genes were down-regulated (adjusted *p* < 0.05), and their repression was more severe (1,135 below twofold cutoff). Genes repressed in the double-CTR mutant included the majority of genes down-regulated by CTR1 mutation alone (*n* = 3,210, **Figure S3D**). The genes most significantly up-regulated in the double-CTR mutant (*n* = 31) were also induced by CTR1 mutation alone, and included genes related to stress responses such as *TENT5C*, *ATF3* and *SARM1* (**Figure S3E**). Overall, gene expression changes in the double-CTR mutant *muR1R2* were highly correlated with those in the CTR1 mutant *muR1* (Pearson correlation; R = 0.87) and in the *muALL* mutant with all three regions mutated (R = 0.88) (**Figure S3F**). The genes down-regulated by CTR1 or double-CTR mutation were not enriched for ones involved in a particular biological process, but tended to be longer than average (**Figure S3G**), consistent with repression being due to defects in elongation.

RNAPII elongation and co-transcriptional splicing of pre-mRNAs are coupled, in part through phosphorylation-dependent mechanisms ^56,57^. To ask if *SUPT5H* mutations affect splicing, we analyzed the frequencies of different alternative splicing (AS) events: skipped exons, alternative 5’ splice sites, alternative 3’ splice sites, mutually exclusive exons, and retained introns (**Figure 3E**). For all classes of AS, the double-CTR mutation produced a higher incidence of significantly changed events than did either CTR mutation alone. The most commonly altered event was exon-skipping, with similar frequencies of up- and down-regulated events, and correspondence between the numbers of genes affected and effects on cell viability and steady-state mRNA levels in each of the mutants (*muR1R2* ≈ *muALL* > *muR1* > *muR2*) (**Figures 3F**, **3G** and **S3H**). More than half the genes with altered splicing patterns in the double-CTR mutant were down-regulated in the RNA-seq analysis (**Figure S3I**). Therefore, preventing CTR2 phosphorylation exacerbated effects of CTR1 mutation on cell proliferation, pre-mRNA splicing and gene expression, consistent with reinforcing functions of the two CTRs.

### Control of pausing by tripartite SPT5 phosphorylation

To ask if CTR1 and CTR2 mutations had similarly reinforcing effects on chromatin distribution of RNAPII, we performed ChIP-seq analysis in the Flag-SPT5-expressing cells after depletion of endogenous dTAG-SPT5 (**Figures 4A-C** and **S4A-C**).

**Figure 4.**
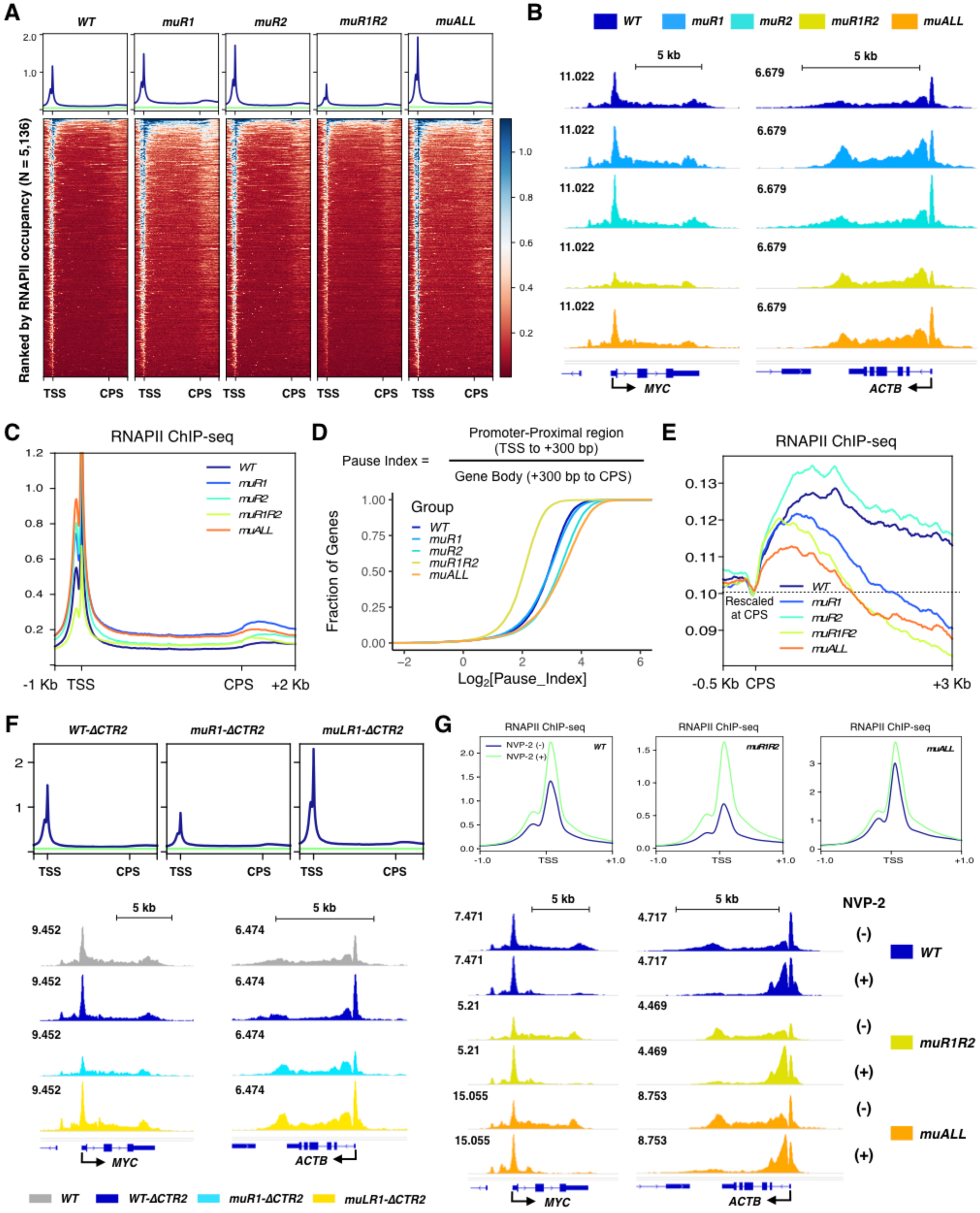
Control of pausing by tripartite SPT5 phosphorylation. **A.** Spike-in normalized metagene plots (top) and heatmaps (bottom) from ChIP-seq analysis of RNAPII (RPB1) in *dTAG-SUPT5H* cells expressing indicated Flag-SPT5 variants after dTAGv-1 treatment (125 nM, 4 hr). Green line: intergenic signal. Data are from two biological replicates (n = 2). **B.** Browser tracks of RNAPII ChIP-seq data at *MYC* and *ACTB* loci. **C.** Merged metagene plot for RNAPII ChIP-seq data. **D.** Empirical cumulative distribution function (ECDF) plot of RNAPII pausing index for genes with high RNAPII occupancy (n = 10,151). Plots represent averaged result from two independent experiments (n = 2). **E.** CPS-centered metagene plot for RNAPII ChIP-seq data. Data were re-scaled at CPS to emphasize changes in the downstream region. **F.** Intergenic signal-normalized metagene plots (top) and browser tracks (bottom) from RNAPII ChIP-seq in *dTAG-SUPT5H* cells expressing indicated CTR2-deleted Flag-SPT5 variants after dTAGv-1 treatment (125 nM, 4 hr). Green line: intergenic signal. Data are from two biological replicates (n = 2). Gene browser track from analysis of full-length, wild-type Flag-SPT5 in (B) is included for comparison. **G.** Spike-in normalized metagene plots (top) and *MYC* and *ACTB* gene browser tracks (bottom) from RNAPII (RPB1) ChIP-seq in cells expressing indicated full-length Flag-SPT5 variants after treatment with DMSO (-) or 250 nM NVP-2 (+) for 1 hr. Data are from two biological replicates (n = 2) for NVP-2-treated *muR1R2* and *muALL*; one (n = 1) for other conditions.

Individually, mutation of CTR1 or CTR2 increased RNAPII occupancy in promoter-proximal regions. In the CTR1 mutant, RNAPII occupancy also increased in gene bodies, resulting in largely unchanged pause index (PI) values (**Figure 4D**). In contrast, CTR2 mutation had little effect on RNAPII occupancy in gene bodies and therefore produced globally increased PIs. Downstream of the CPS, the CTR1 mutation alone caused a more precipitous drop in RNAPII occupancy compared to wild-type SPT5, as previously reported ^30^, and this pattern was not affected by additional mutation of CTR2 (**Figure 4E**). Unexpectedly, however, blocking phosphorylation of both CTRs caused a large, global decrease in promoter-proximal pausing (**Figures 4A**-**D**, **S4B** and **S4C**).

Reduced promoter-proximal occupancy by RNAPII in cells expressing the *muR1R2* variant might be explained by decreased initiation, diminished pausing, or both. To distinguish among these possibilities, we performed ChIP-seq analysis in cells expressing the *muALL* variant. Addition of the KOWx-4/5 linker mutations overrode the effect of dual-CTR mutation on pausing and restored RNAPII occupancy in the promoter-proximal region (**Figure 4A**-**D**). Analysis of the allelic series containing internal deletion rather than point mutations of CTR2 yielded similar results (**Figures 4F** and **S4D-G**): promoter-proximal RNAPII occupancy was diminished by an SPT5 variant in which neither CTR could be phosphorylated (*muR1τιCTR2*) and restored by additional mutation of the KOWx-4/5 linker (*muLR1τιCTR2*). The KOWx-4/5 linker binds RNA and is positioned over the RNA exit channel in the paused RNAPII elongation complex ^58^, and mutations that prevent its phosphorylation impede pause release ^30,47^. These results therefore suggest that diminished promoter-proximal RNAPII occupancy in the double-CTR mutant was primarily due to accelerated pause release, and could be restored by preventing KOWx-4/5 linker phosphorylation.

The SPT5 *muALL* variant is devoid of S/T-P sites matching the minimal consensus motif for phosphorylation by CDKs. Cells expressing this variant nonetheless had an RNAPII distribution similar to that in wild-type cells, with peaks of occupancy consistent with pausing in promoter-proximal and 3’ regions (**Figures 4B** and **S4B**). To ask if RNAPII elongation remained responsive to CDK9 inhibition, we treated cells expressing the *WT*, *muR1R2* or *muALL* variants with NVP-2 for 1 hr prior to ChIP-seq analysis. The qualitative response to CDK9 inhibition was similar in all three cases, with increased RNAPII accumulation in the promoter-proximal region and diminished occupancy in the gene body and termination zone (**Figure 4G**). The effect of NVP-2 near the TSS was greatest in cells expressing the *muR1R2* variant and least for the *muALL* variant, indicating that the CDK9 inhibitor phenocopied KOWx-4/5 linker mutations by impeding release of promoter-proximally paused RNAPII in the double-CTR mutant. Nonetheless, elongation remained sensitive to NVP-2 even when SPT5 lacked all known CDK phosphorylation sites, suggesting that other substrates, such as NELF, the PAF complex, SPT6 or RNAPII itself ^25^, are capable of conferring regulation by CDK9.

### Combinatorial effects of CTR1 and CTR2 mutations on nascent transcription

To try to reconcile the additive effects of CTR mutations on steady-state mRNA levels and pre-mRNA splicing with their impacts on RNAPII distribution, we analyzed nascent RNA synthesis in cells expressing wild-type or mutant Flag-SPT5 by transient transcriptome sequencing with chemical fragmentation of RNA (TT_chem_-seq) ^59^. Consistent with the RNA-seq results (**Figures 3C** and **3D**), preventing phosphorylation of CTR1 alone was generally repressive of nascent transcription, with significant down-regulation of 679 transcripts and up-regulation of 44 (**Figures 5A** and **S5A**, adjusted *p* < 0.05), whereas mutating CTR2 alone, which had no significant gene-specific effects (adjusted *p* < 0.05) (**Figure 5A**), modestly increased nascent RNA synthesis globally, most prominently on highly expressed genes (**Figures S5A** and **S5B**). When combined with mutation of CTR1, however, the CTR2 mutation had a dramatic and largely positive effect on nascent transcription; globally, transcriptional output was restored to near wild-type levels (**Figure S5B**), and significantly up-regulated transcripts (**Figures 5A** and **5B**, *n* = 1,077, adjusted *p* < 0.05) now outnumbered down-regulated ones (*n* = 708, adjusted *p* < 0.05). The transcripts most strongly repressed by the CTR1 mutation, however, remained so in the double-CTR mutant (**Figure 5C**). Addition of KOWx-4/5 linker mutations dampened the gene-specific effects of the double-CTR mutation—both up- and down-regulated transcripts were fewer in number (**Figures 5A** and **S5A**)—but had only a modest impact on metagene plots of nascent transcription (**Figures 5B** and **S5B**).

**Figure 5.**
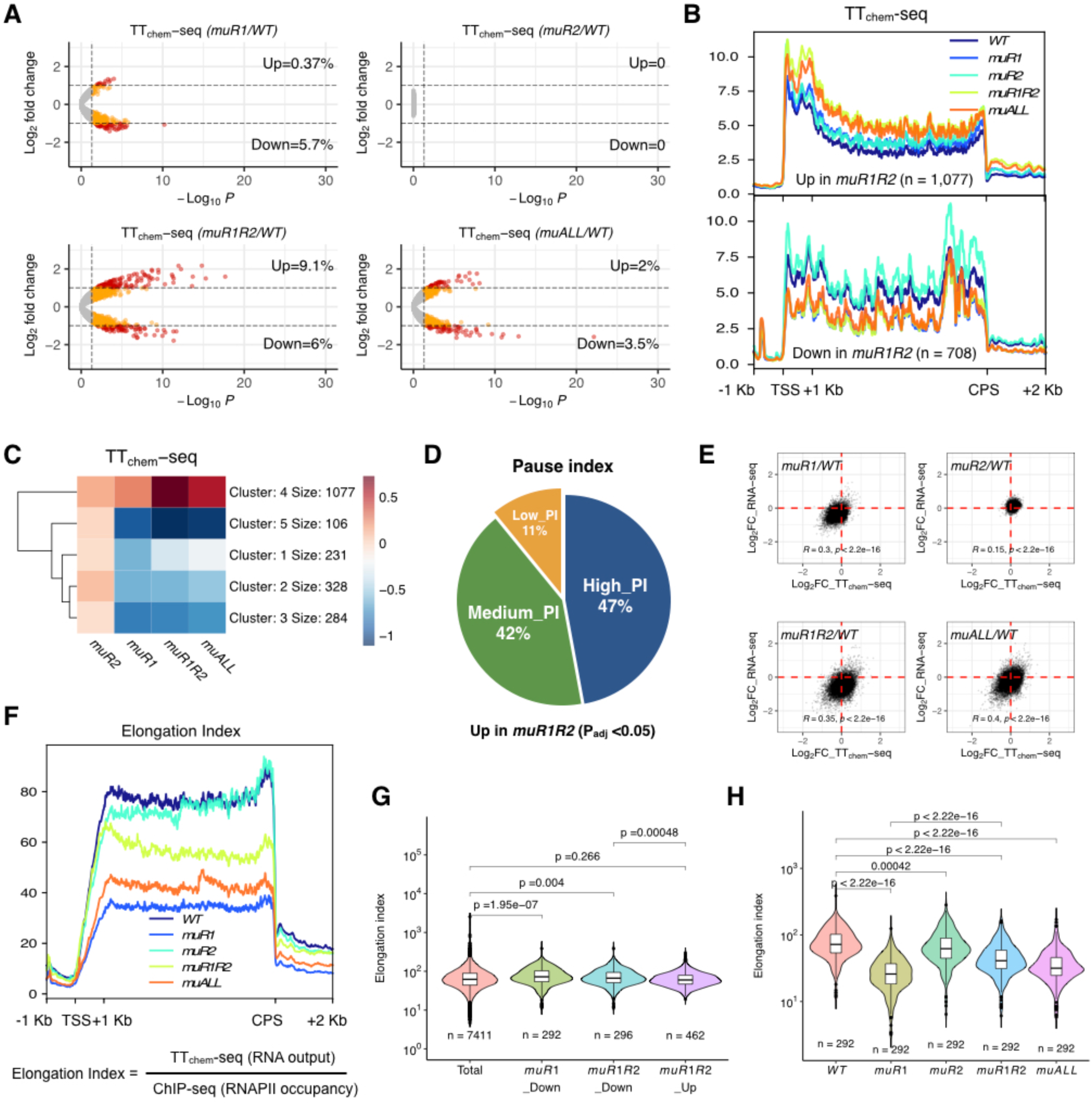
Combinatorial effects of CTR1 and CTR2 mutations on nascent transcription. **A.** Volcano plots of spike-in TT_chem_-seq data in cells expressing indicated Flag-SPT5 variants after dTAGv-1 treatment (125 nM, 4 hr). Colored dots represent significantly changed genes in differential gene analysis (Wald test, adjusted p < 0.05, n = 11,841). Red dots indicate genes with fold-change > 2. Data are from two or three biological replicates (n = 2 for *muR1* and *muALL*; n = 3 for *WT*, *muR2*, *muR1R2*). **B.** Merged metagene plots (sense strand) of TT_chem_-seq data. The genes significantly changed by double-CTR mutation (adjusted p < 0.05) were selected (n = 1,077, Up; n = 708, Down). **C.** Heat map of TT_chem_-seq data for genes significantly changed by CTR1 and double-CTR mutation (adjusted p < 0.05), aggregated into 5 clusters by k means clustering. **D.** Pie chart of nascent transcripts up-regulated in the double-CTR mutant. The corresponding genes were divided by pause index (PI) into groups with high, medium, and low pausing propensity. **E.** Scatterplots of RNA-seq and TT_chem_-seq data. The log2 transformed fold changes in TT_chem_-seq and RNA-seq are plotted on *x*- and *y*-axes, respectively. Pearson correlation R and p values are indicated. **F.** Merged metagene plots (sense strand) for the elongation index (EI) across non-overlapping genes longer than 1 kb (n = 7,448). EI was defined as indicated. **G.** Violin plots of EI (*WT*) for the indicated gene sets. Wilcoxon signed rank test, gene number (n) and p-values are as indicated. **H.** Violin plots of EI in indicated SPT5 variants for genes down-regulated by CTR1 mutation. Wilcoxon signed rank test, gene number (n) and p-values are indicated.

We suspected dampening of nascent transcription by the KOWx-4/5 linker mutation might reflect restored pause regulation. Consistent with this explanation, when we sorted genes according to PI in wild-type cells, the up-regulated gene set in the double-CTR mutant was enriched for moderately (Log_2_PI ≥ 2.0 and < 3.0) and highly pause-regulated genes (Log_2_PI ≥ 3.0), which accounted for ∼89% of the up-regulated transcripts, compared to ∼69% of all genes (**Figure 5D**). In addition, genes involved in transcription initiation or elongation and DNA damage responses were overrepresented (**Figure S5C**). There was only modest correlation between changes in nascent and steady-state RNA levels in the various mutants (**Figure 5E**); many transcripts enriched in TT_chem_-seq analysis of the double-CTR mutant were depleted in RNA-seq analysis (lower right quadrant), suggesting they were unstable.

Compared to the CTR1 mutant, the double-CTR mutant variant of SPT5 supported higher transcriptional output with lower gene-body occupancy by RNAPII (**Figure S5D**), suggesting an increased elongation rate. To explore this possibility, we calculated the elongation index (EI) ^60^ by dividing TT_chem_-seq by ChIP-seq signals for each SPT5 mutant (**Figure 5F**). By itself, CTR1 mutation decreased EIs, consistent with slowed elongation, whereas mutating CTR2 alone had little effect. The double-CTR mutation resulted in EIs intermediate between those of wild-type SPT5 and the CTR1 mutant, suggesting that 1) in the absence of CTR1 phosphorylation, unopposed CTR2 phosphorylation acts as a brake on RNAPII elongation; and 2) CTR1 phosphorylation is needed to attain maximal elongation speeds, even when CTR2-dependent braking is disabled. Consistent with this model, the gene sets with diminished nascent transcription in both CTR1 and double-CTR mutants had significantly higher EIs in wild-type cells (**Figure 5G**), suggesting that faster-elongated transcripts were more sensitive to CTR1 phosphorylation. Moreover, for transcripts downregulated in the CTR1 mutant, simultaneous mutation of CTR2 produced a highly significant increase in EI (**Figure 5H**), consistent with opposing effects of CTR1 and CTR2 phosphorylation on elongation rate.

### Dichotomous regulation of transcription termination by the SPT5 CTRs

Closer inspection of the genes most strongly up-regulated in the double-CTR mutant revealed that, in most cases, increased TT_chem_-seq signals resulted from read-through transcription originating in more heavily transcribed upstream genes. This was evident in individual gene browser tracks, and validated by RT-qPCR (**Figures S6A** and **S6B**). At these poorly transcribed downstream genes, the CTR1 mutation alone gave rise to mildly increased nascent transcription, which was enhanced in the double-CTR mutant (**Figure S6C**). Of the upstream genes that gave rise to read-through transcription, 41.4% and 62.2% were repressed at the steady-state mRNA level by CTR1 and double-CTR mutations, respectively (**Figure S6D**), possibly owing to defects in 3’-end maturation.

To ask if transcriptional read-through occurred more broadly in the double-CTR mutant, we analyzed nascent RNA signals genome-wide in the 5 kilobases (kb) downstream of the CPS. Mutating both CTRs increased TT_chem_-seq signals downstream of the CPS at a large number of genes (*n* = 1,307; signal > 1 downstream of the CPS in *muR1R2*; fold-change ≥ 1.5), whereas mutating CTR1 alone increased read-through transcription more modestly **(Figures 6A** and **S6E)**. Our genome-wide analysis of ChIP-seq data had revealed abrupt decreases in RNAPII occupancy downstream of the CPS in either the CTR1 or double-CTR mutant (**Figure 4E**), suggesting faster termination, and appeared to corroborate a previous study that analyzed a similar CTR1-mutant variant ^30^. To probe this seeming discrepancy, we queried TT_chem_-seq data for genes with reduced nascent RNA signal in the 5 kb downstream of the CPS, using a similar threshold (signal > 1 downstream of the CPS in *WT*; fold-change ≤ 0.67). By these criteria, 866 genes appeared to undergo faster termination in the CTR1 or double-CTR mutant **(Figures 6B** and **S6F**). For both sets of genes, there was agreement between TT_chem_-seq and ChIP-seq analyses; both nascent transcription and RNAPII occupancy remained elevated downstream of the CPS in the double-CTR mutant at genes prone to read-through, but both decreased at genes undergoing faster termination (**Figures S6G**). We saw a similar dichotomy in plots of EI in the 5-kb window downstream of the CPS for the read-through (*muR1R2* > *WT > muR1*) (**Figure 6C**) and fast-terminating (*WT* > *muR1R2 > muR1*) gene sets (**Figure 6D**), suggesting faster elongation through the termination zone in the former, but slower elongation in the latter, when all CTR phosphorylation was prevented.

**Figure 6.**
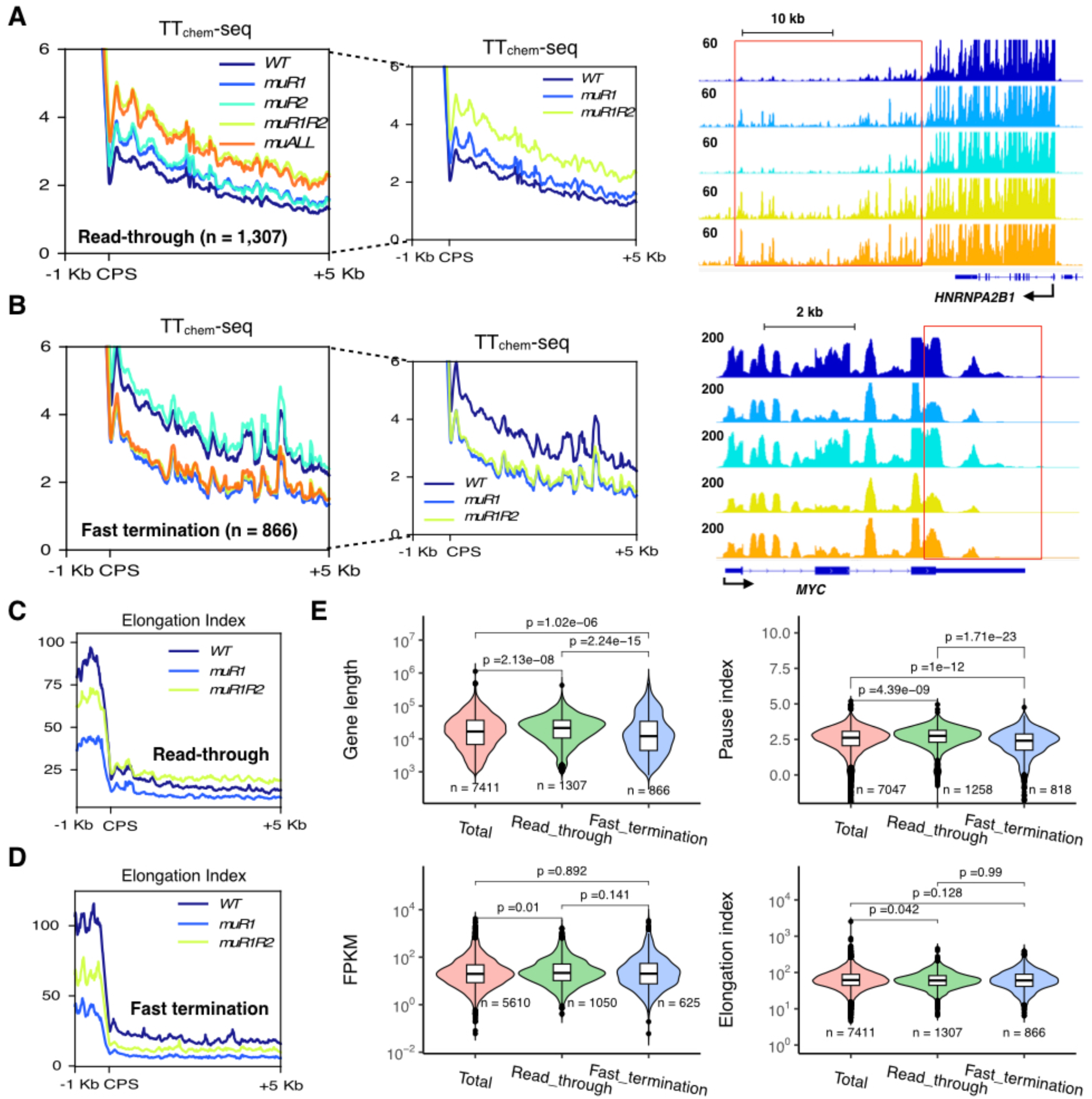
Dichotomous regulation of transcription termination by the SPT5 CTRs. **A.** CPS-centered metagene plots (left and middle) and browser tracks (right) of TT_chem_-seq data for read-through genes in *muR1R2* (*n* = 1,307; signal > 1 in *muR1R2*; fold-change ≥ 1.5). Middle plot shows same data as at left, with *muR2* and *muALL* data removed to allow easier comparison. **B.** CPS centered metagene plots (left and middle) and browser tracks (right) of TT_chem_-seq data for fast-termination genes in *muR1R2* (*n* = 866; signal > 1 in *WT*; fold-change ≤ 0.67). Middle plot shows same data as at left, with *muR2* and *muALL* data removed to allow easier comparison. **C.** CPS centered metagene plots (sense strand) of EI of read-through genes in *muR1R2* (*n* = 1,307). **D.** CPS centered metagene plots (sense strand) of EI of fast-termination genes in *muR1R2* (*n* = 866). **E.** Violin plots of gene length, PI, expression level (FPKM), and EI for the indicated gene sets (all parameters are from *WT*). Wilcoxon signed rank test, gene number (n) and p-value are as indicated.

To identify gene characteristics associated with different termination behaviors, we compared gene length, RNA expression level, PI and EI (all parameters measured in wild-type cells) for the read-through and fast-terminating gene sets (**Figure 6E**). This analysis revealed that longer genes with higher PIs were more prone to read-through, whereas shorter, less paused genes were more prone to faster termination. In addition, among genes with delayed termination, there was a slight but significant bias towards higher expression levels and slower elongation. Both gene sets overlapped extensively with genes down-regulated in RNA-seq analysis of the double-CTR mutant, but there was little overlap with the genes that displayed aberrant splicing (**Figure S6H**). Together, alterations in termination and splicing can account for ∼22% of genes with decreased steady-state mRNA levels in cells expressing the double-CTR mutant variant of SPT5.

### CTR1 and CTR2: accelerator and brake on RNAPII elongation

The effects of CTR1 and CTR2 mutations on EI suggested complex regulation of RNAPII elongation velocity by CTR phosphorylation. To measure elongation rates directly in cells dependent on different SPT5 variants, we reversibly blocked RNAPII in the promoter-proximal region by a 3-hr treatment with the CDK inhibitor 5,6-dichloro-1-β-D-ribofuranosyl-benzimidazole (DRB), and monitored progression of the transcribing RNAPII wavefront by CUT&RUN at 0, 3 and 5 min after DRB washout (**Figure 7A**). We observed slowing of RNAPII transit in the CTR1 single mutant, which was partially reversed in the double-CTR mutant, on individual genes (**Figures 7B** and **S7A**), and in metagene analysis of non-overlapping, highly expressed genes > 10 kb in length (**Figure 7C**). In the presence of an intact CTR1, the CTR2 mutation alone had no significant effect on elongation rates (**Figures S7B** and **S7C**). The additional mutations of the KOWx-4/5 linker in the *muALL* variant further impeded progression of transcribing RNAPII (**Figures 7B** and **7C**).

**Figure 7.**
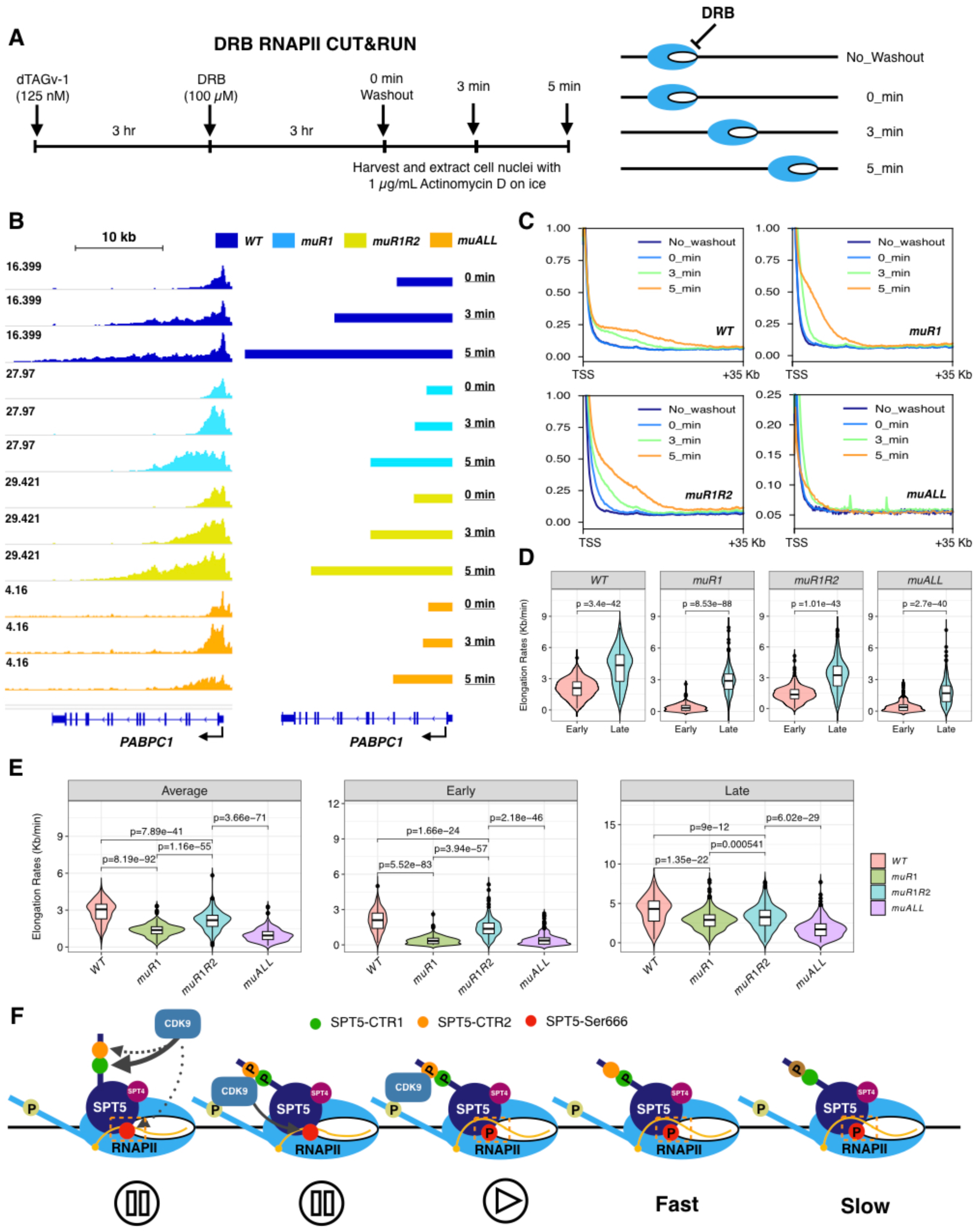
CTR1 and CTR2: accelerator and brake on RNAPII elongation. **A.** Schematic of the DRB-washout RNAPII (RPB1)-CUT&RUN protocol. **B.** Browser tracks (left) and wave tracks (right) of RNAPII CUT&RUN data at the *PABPC1* gene. Data are from two biological replicates (n = 2). **C.** Merged metagene plots of RNAPII CUT&RUN data at indicated time points for non-overlapping, highly expressed genes > 10 kb in length (n = 4,152). Data are from two biological replicates (n = 2). **D.** Violin plots of RNAPII elongation rates measured at early and late intervals after release from DRB on non-overlapping, highly expressed genes longer than 30 kb (n = 255). Paired t-test, p-value is as indicated. **E.** Violin plots of RNAPII elongation rates measured at early and late intervals after release from DRB on non-overlapping, highly expressed genes > 30 kb in length (n = 255). Paired t-test, p-values are indicated. **F.** A model summarizing the proposed functions of SPT5 phosphorylations in CTR1, CTR2, and the KOWx-4/5 linker in RNAPII transcription elongation. DSIF (a complex of SPT5 and SPT4) impedes elongation by RNAPII in the promoter-proximal region when the RNA clamp (KOWx-4/5 linker) is unphosphorylated. CDK9/CycT1 phosphorylates CTR1, which can potentiate high-speed RNAPII elongation, while DSIF-RNAPII complex remains paused. CDK9/CycT1 also phosphorylates CTR2, which can slow RNAPII elongation when CTR1 is unphosphorylated. RNAPII with associated DSIF becomes fully competent for elongation once the KOWx-4/5 linker is phosphorylated.

The farther advance of the RNAPII wavefront in the double-CTR mutant might reflect faster release from the promoter-proximal pause, an increased intrinsic rate of elongation, or both. To parse contributions of these two effects, we derived elongation rates from CUT&RUN data in the two time intervals after DRB washout, 0-3 min and 3-5 min, for a subset of non-overlapping, highly expressed genes > 30 kb in length (**Table S1**). For all four *SUPT5H* genotypes—*WT*, *muR1*, *muR1R2* and *muALL*—elongation was slower during the first interval and sped up between 3 and 5 min after release from the DRB block (**Figure 7D**). Direct comparison of the variants revealed the greatest differences during the first 3 min after DRB washout; between 3 and 5 min, there were smaller but still significant differences in RNAPII velocity, in the order *WT* > *muR1R2* > *muR1 > muALL* (**Figures 7E**). Therefore, both positive and negative regulation of RNAPII speed by SPT5 CTR phosphorylation persisted well into the elongation phase. Addition of the KOWx-4/5 linker mutation also slowed elongation, relative to the double-CTR mutant, in both time intervals, suggesting that phosphorylation in the RNA-binding loop of SPT5 can override a lack of CTR phosphorylation to promote elongation in gene bodies as well as the promoter-proximal region.

## DISCUSSION

Precise regulation of transcript elongation is an ancient mode of gene expression control ^11^. The prokaryotic ortholog of SPT5, NusG ^61^, has pro- or anti-pausing effects on transcription in different bacterial species. Eukaryotic SPT5 likewise acts as either a negative or positive regulator of RNAPII elongation, depending on its phosphorylation state ^31,62^. Recent work has revealed that phosphorylation by P-TEFb does not simply flip a binary switch on SPT5; besides the repetitive carboxy-terminal motifs through which CDK9 seemed to promote elongation ^34–36^, human SPT5 contains sites in the KOWx-4/5 linker that were phosphorylated by CDK9 during activation of the paused elongation complex in vitro ^25^, mutation of which impeded pause release in cells ^30,47^. Importantly, while the same kinase phosphorylates the KOWx-4/5 linker and CTR1 of human SPT5, different phosphatases remove these marks ^15,44,46^, potentially explaining their different distributions on chromatin ^15^ and allowing them to regulate different steps in the transcription cycle ^14^.

Phosphorylation of SPT5 by P-TEFb was known to promote elongation, but here we reveal additional complexity. Human SPT5 contains two carboxy-terminal, S/T-P-rich regions, which exert opposite effects on RNAPII elongation speed—acceleration by pCTR1 and deceleration by pCTR2 (**Figure 7F**). Loss of either input had mild or moderate effects on steady-state mRNA levels and cell proliferation, but loss of both severely compromised gene expression and viability. Two potential explanations for these combinatorial effects emerged from transcriptomic analyses of the double-CTR mutant variant—increased frequency of aberrant splicing events and altered termination efficiency, both of which correlated with diminished steady-state mRNA levels. Skipping of a given exon was ∼equally likely to be increased or decreased by the double-CTR mutation, as predicted if the primary defect is in regulating elongation speed; slowed elongation can favor either inclusion or exclusion of alternative exons ^63^. Interestingly, mutating both CTRs also had divergent effects on termination—transcriptional read-through or termination nearer the CPS—at different genes. We infer that the loss of control by SPT5 phosphorylation, rather than a changed rate of RNAPII elongation per se, gave rise to the transcriptional derangements we observed. For individual genes, the two mechanisms appear to be mutually exclusive—the sets of genes that displayed aberrant splicing or termination patterns were largely non-overlapping—and would thus be expected to have additive effects on gene expression and cell viability.

The slowing of RNAPII transcription by CTR1 mutation preferentially affected early elongation, resembling the effects of CDK9 inhibition in both human and yeast systems ^35,64^. This indicates a role in regulating the onset of rapid elongation at or, more likely, soon after pause release; RNAPII occupancy was increased in the promoter-proximal region and throughout gene bodies, consistent with a generally reduced rate of elongation. By itself, CTR2 mutation increased pausing without affecting RNAPII gene-body occupancy or elongation rate. Mutating both CTRs globally attenuated pausing, however, even as it partially rectified elongation rates. Preventing KOWx-4/5 linker phosphorylation or inhibiting CDK9 overrode this pause attenuation, suggesting that the inability of CDK9 to phosphorylate the CTRs led to premature or inappropriate pause release that nonetheless remained dependent on linker phosphorylation.

CTR1, CTR2 and the KOWx-4/5 linker are all CDK9 substrates ^25,31–33,39,51^, phosphorylation of which is NVP-2-sensitive in human cells (^38^, this report). Therefore, a possible explanation for our results is that multi-site phosphorylation of the SPT5 CTRs serves a timing function, perhaps by competing with the KOWx-4/5 linker for limiting CDK9 and thus ensuring that pause release occurs only after a threshold level of CTR phosphorylation is reached (**Figure 7F**). Consistent with this model, pCTR1 peaked in the promoter-proximal region in unperturbed HCT116 cells, whereas pSer666 was depleted over the pause before increasing in the downstream region ^38^.

Our results may help explain those of a previous genetic dissection of zebrafish SPT5 ^52^, in which removal of either CTR1 or CTR2 had little or no effect, while a truncation removing both CTRs severely disrupted embryonic development, perhaps analogous to the synthetic-lethal effect of dual-CTR mutation we observed in human cells. Carboxy-terminal truncation of zebrafish SPT5 specifically impaired its ability to restrain elongation; heat shock promoter-driven GFP expression was de-repressed, in the absence of a heat shock, by expression of the truncated SPT5 variant. These effects make sense if the CTRs of zebrafish SPT5 help prevent precocious pause release, as they appear to do in HCT116 cells.

We note that, in a recent mutational analysis of SPT5 also performed in HCT116 cells, pausing persisted in a CTR1 mutant with T→A substitutions of the seven sites phosphorylated by CDK9, even though sequences carboxy-terminal to CTR1, including CTR2, had been removed ^47^. This discrepancy cannot be explained simply by a difference between CTR2 excision and point mutation, because the internal deletion of CTR2 analyzed here recapitulated effects of the substitutions on RNAPII distribution, including effects on pausing (**Figures 4F** and **S4D-G**). Possible explanations include 1) a previously undetected function of residues located distal to CTR2, such as the KOW6-7 domain (**Figure 2A**), which when mutated caused defects in neurodevelopment in zebrafish ^65^; or 2) a context-dependent effect of CTR1 phosphorylation—or lack thereof—when the natural carboxyl terminus of the protein is missing. Interestingly, Bentley and co-workers observed pause attenuation when they introduced T→E substitutions in CTR1 of a truncated, CTR2-less SPT5 ^47^, perhaps consistent with competition between CTR1 and the KOWx-4/5 linker for limiting CDK9.

Phosphorylation-site mutations of CTR2 did not by themselves produce obvious phenotypes or globally alter elongation rates unless CTR1 was also mutated, but two observations suggest a role for CTR2 phosphorylation in unperturbed transcription cycles. First, the isoform phosphorylated on CTR2 is a significant fraction of chromatin-associated SPT5 in cells expressing wild-type SPT5 either from the endogenous locus or ectopically. Second, mutations preventing CTR2 phosphorylation mildly enhanced nascent transcription and steady-state mRNA levels, and increased RNAPII occupancy in the promoter-proximal region, even when sites in CTR1 and the KOWx-4/5 linker remained intact. Taken together, our results suggest that human SPT5 contains three distinct regions phosphorylated during the RNAPII transcription cycle: the KOWx-4/5 linker, which in unmodified form restrains pause release irrespective of CTR phosphorylation status; CTR1, phosphorylation of which accelerates elongation after release from the pause; and CTR2, which may act as a throttle or brake to limit elongation speed and prevent runaways—transcription through natural stop signals, or despite errors in co-transcriptional RNA processing—that might otherwise result from unopposed CTR1 phosphorylation.

## LIMITATIONS OF THE STUDY

We performed all analyses in cells derived from a single cancer cell line—a practical necessity given the chemical genetic strategy used and the number of SPT5 mutant variants analyzed. Given the conserved function of SPT5 in transcription, however, we believe that significant differences between cell types are unlikely. Our initial detection of CTR2 phosphorylation was serendipitous; the increased phosphorylation of CTR2 when CTR1 phosphorylation was prevented by mutation uncovered the cryptic cross-reactivity of anti-pSer666 with pCTR2. We have not determined which of the 22 S/T-P sites in CTR2 cross-react with the antibody when phosphorylated, but the increased signal upon CTR1 mutation suggests that the relevant epitope might not be heavily phosphorylated under physiologic conditions, even though CTR2 phosphorylation clearly occurs, as indicated by the phosphatase-sensitive electrophoretic mobility shift. The genome-wide acceleration of elongation when CTR2 phosphorylation was blocked was only detected in the absence of CTR1 phosphorylation. Therefore, further studies will be needed to determine whether the braking function of CTR2 limits elongation rates under physiologic conditions, for example on specific genes or within defined gene regions. Conversely, slowing of elongation when CTR1 phosphorylation is prevented might reflect direct effects of pCTR1 on the RNAPII elongation complex, or an indirect effect due to the increased CTR2 phosphorylation caused by this mutation. Finally, some of the phenotypic effects we observed might stem from the mutations themselves, rather than any effect on CTR phosphorylation; SPT5 was recently shown to undergo phase separation, dependent on intrinsically disordered regions in the CTRs, but not strictly dependent on their phosphorylation ^66^.

## ACKNOWLEDGMENTS

We thank P.K. Parua (Albert Einstein College of Medicine) for helpful discussions during the planning stages of the study, M.J. Gamble (Albert Einstein College of Medicine) for helpful discussions and critical review of the manuscript, and N.S. Gray (Stanford University) for providing NVP-2. This work was supported by National Institutes of Health grant R35 GM127289 to R.P.F. Next-generation sequencing was supported in part by grant P30 CA196521 to the Tisch Cancer Institute. Data analysis used the high performance computing resource at Mount Sinai supported by CTSA grant UL1TR004419.

## AUTHOR CONTRIBUTIONS

R.S. and R.P.F. conceived and planned the study. R.S. conducted all experiments and performed all data analysis. R.S. and R.P.F. wrote the manuscript. R.P.F. conceived the project and acquired funding.

## SUPPLEMENTAL INFORMATION

**Figure S1.**
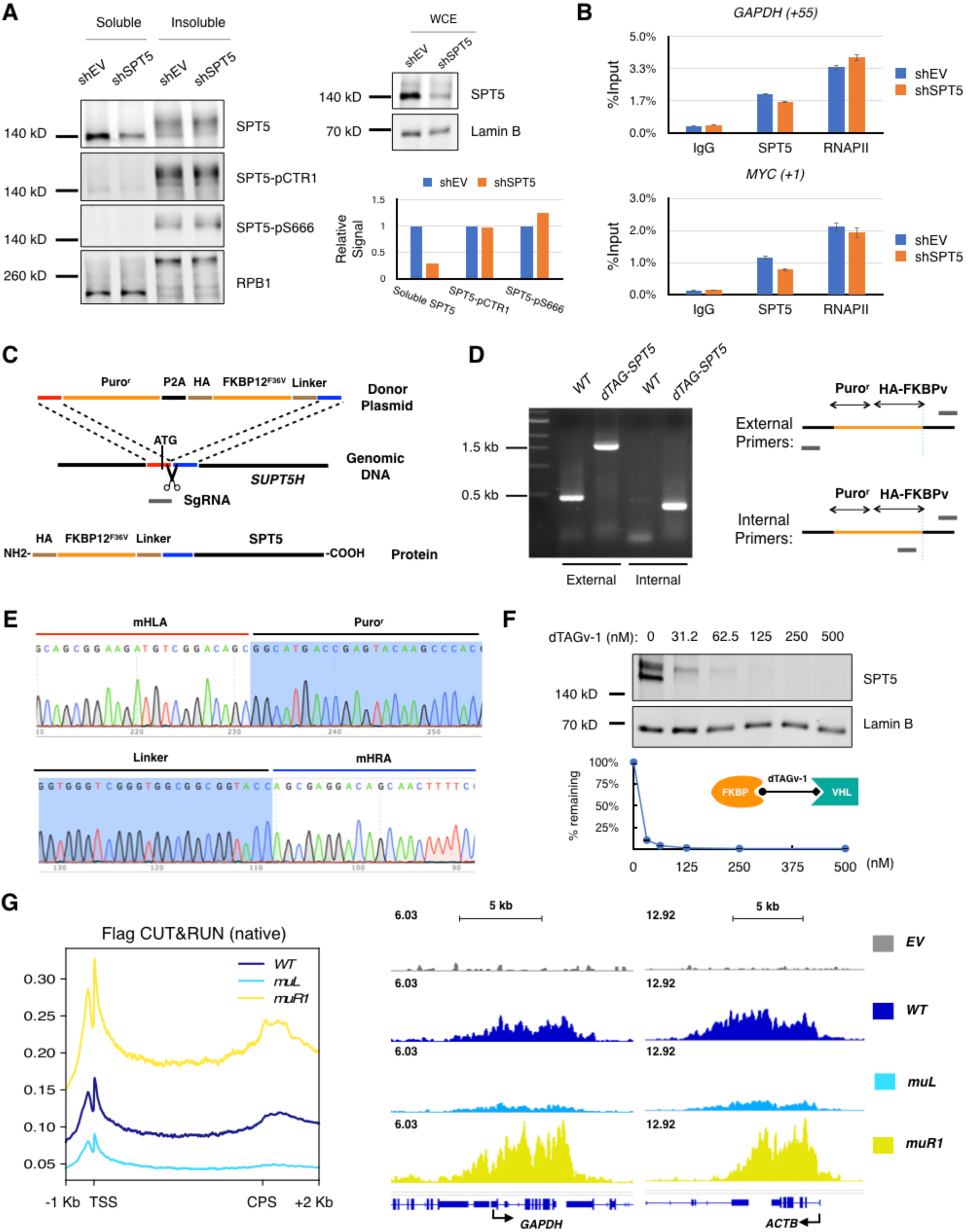
A chemical genetic system for functional dissection of SPT5. Related to Figure 1. **A.** SPT5 depletion by RNAi. Left: HCT116 cells were infected with lentivirus expressing shRNA targeting *SUPT5H* (shSPT5) or empty vector (shEV), fractionated into soluble and insoluble (chromatin) fractions, and subjected to immunoblot analysis with indicated antibodies. Top right: Immunoblot of whole-cell extract (WCE) of HCT116 cells expressing shRNA targeting *SUPT5H* (shSPT5) or empty vector (shEV), probed with antibodies for total SPT5 and Lamin B. Bottom right: Quantification of data in immunoblot at left. **B.** ChIP-qPCR analysis of SPT5 and RNAPII (RPB1) occupancy in *GAPDH* and *MYC* promoter regions in cells expressing shRNA targeting *SUPT5H* (shSPT5) or empty vector (shEV). Data are from two independent experiments (n = 2); error bars indicate mean ± SD. **C.** Schematic of *dTAG-SUPT5H* genome editing strategy. Puro^r^: puromycin resistance gene; P2A: 2A peptide sequence; HA: human influenza hemagglutinin tag; FKBP12^F36V^; degradation tag (dTAG); Linker sequence: (GGGG)S/T x3. **D.** PCR validation of successful genome editing. **E.** Sequencing of edited *dTAG-SUPT5H* loci. The left and right microhomology arms (mHLA and mHRA, respectively) were duplicated in frame with the SPT5 coding sequence. **F.** Dose response of dTAG-SPT5 degradation by dTAGv-1. Top: *dTAG-SUPT5H* cells were incubated for 4 hr with indicated concentrations of dTAGv-1 and whole-cell extracts were prepared and analyzed by immunoblotting for SPT5 and Lamin B. Bottom: Quantification of immunoblot in top panel. **G.** Spike-in normalized metagene plot (left) and browser tracks (right) of Flag CUT&RUN in *dTAG-SUPT5H* cells expressing indicated Flag-SPT5 variants 4 hr after addition of 125 nM dTAGv-1. Data are from two biological replicates (n = 2).

**Figure S2.**
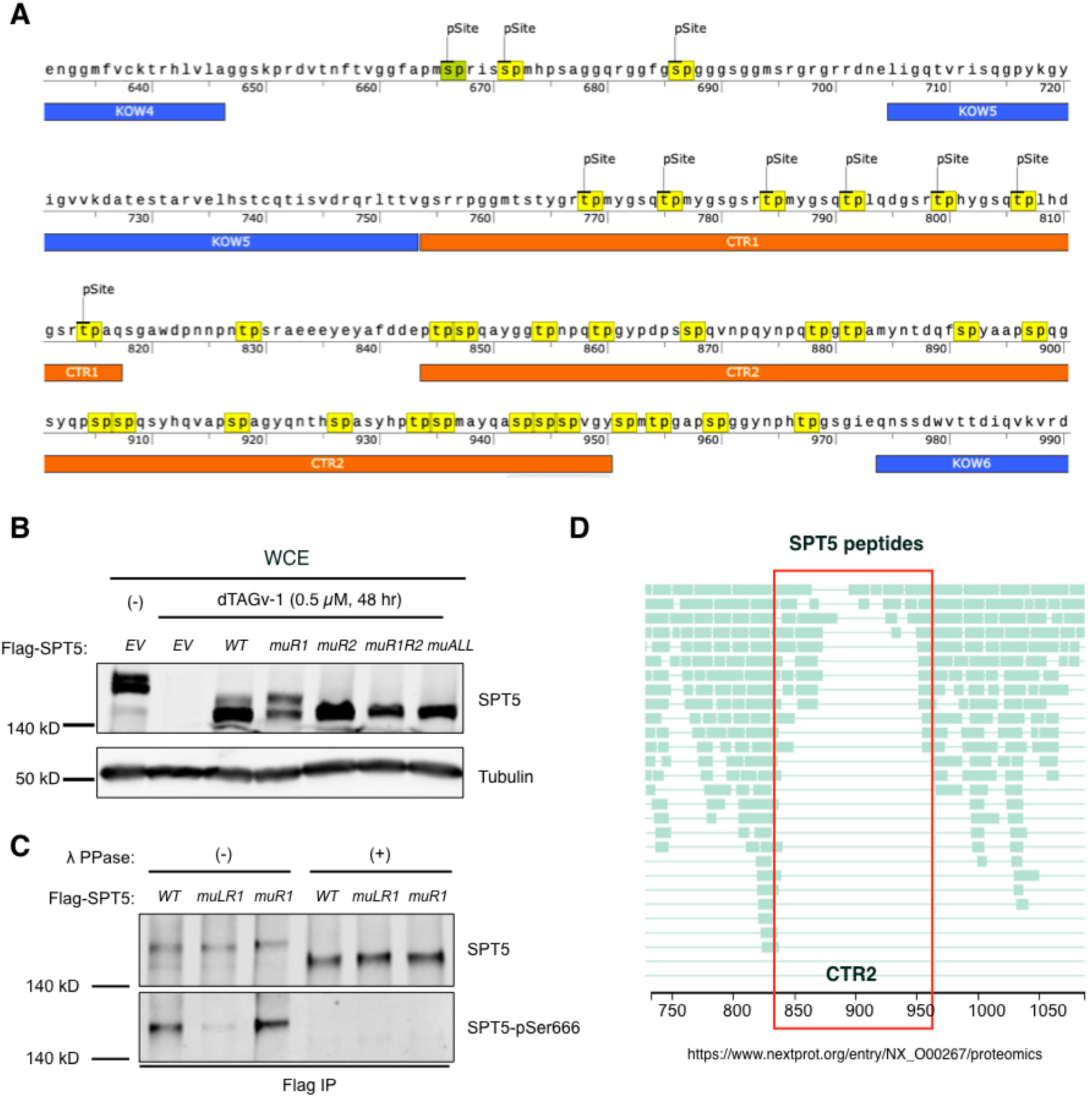
SPT5 CTR2 is phosphorylated in human cells. Related to Figure 2. **A.** Sequence of amino acid residues 631-990 of human SPT5 with potential sites of phosphorylation by CDKs (S/T-P) highlighted in yellow. **B.** Effect of mutations on SPT5 electrophoretic mobility. Whole-cell extract (WCE) was prepared after 48 hr treatment with DMSO (-) or 500 nM dTAGv-1 from *dTAG-SUPT5H* cells expressing indicated Flag-SPT5 variants or empty vector (*EV*) and subjected to immunoblot analysis for total SPT5 and tubulin. **C.** Anti-Flag immunoprecipitates from *dTAG-SUPT5H* cells expressing indicated Flag-SPT5 variants and treated with 125 nM dTAGv-1 for 4 hr were mock-treated (-) or treated (+) with lambda phosphatase (λ PPase) and subjected to immunoblot analysis for total SPT5 and SPT5 phosphorylated at Ser666. **D.** Mass-spectrometric coverage of human SPT5 in public databases. Position of CTR2 is demarcated by red box.

**Figure S3.**
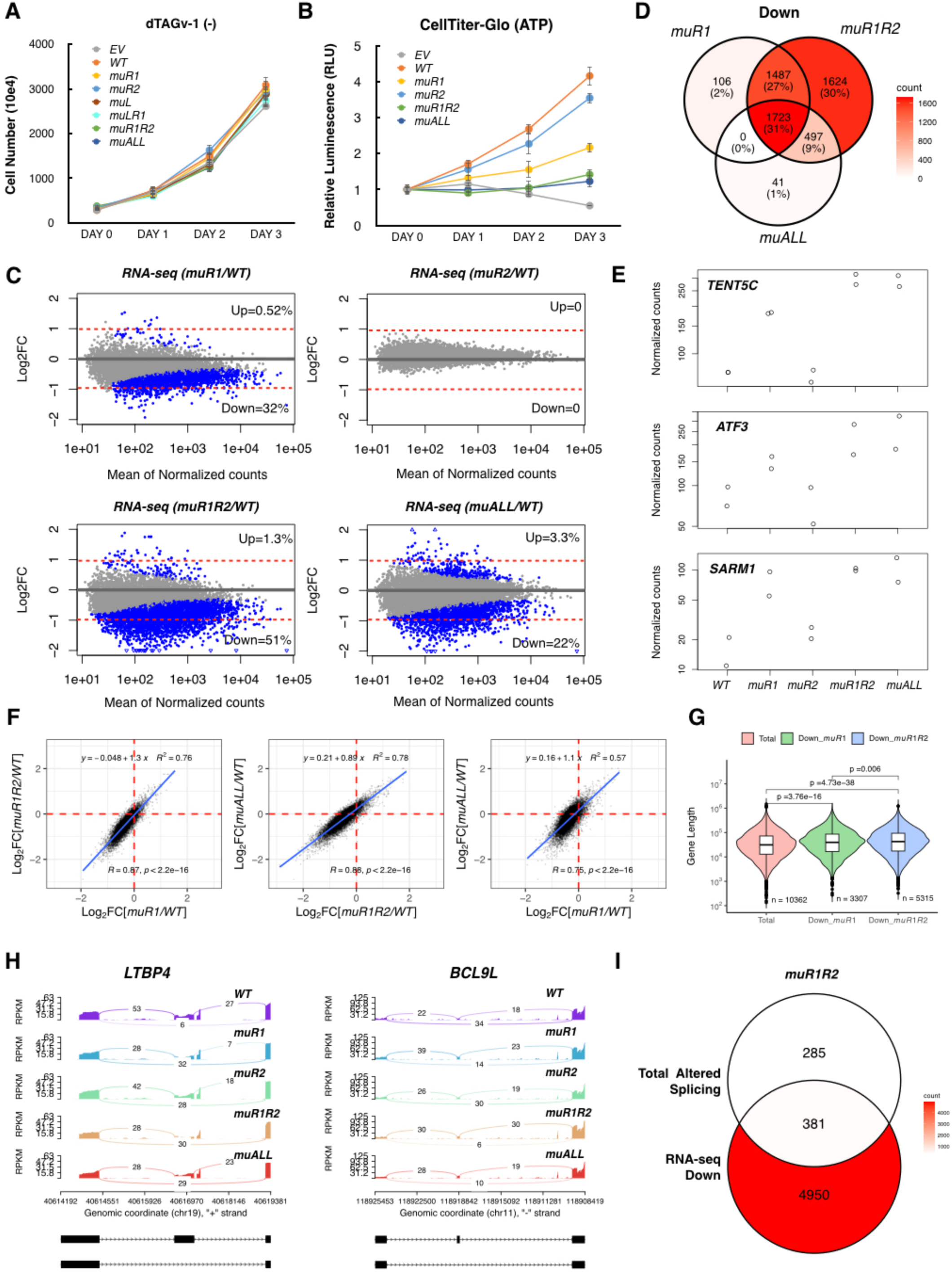
Reinforcing functions of CTR1 and CTR2 in gene expression and cell viability. Related to Figure 3. **A.** Growth curves of *dTAG-SUPT5H* cells expressing indicated Flag-SPT5 variants in the absence of dTAGv-1 treatment. Data are from three independent experiments (n = 3); error bars indicate mean ± SD. **B.** ATP levels measured by CellTiter-Glo assay in *dTAG-SUPT5H* cells expressing indicated Flag-SPT5 variants after indicated times of treatment with 500 nM dTAGv-1. Data are from four independent experiments (n = 4); error bars indicate mean ± SD. **C.** MA plots of RNA-seq data. The *y-*axis shows log2 transformed fold changes and *x-*axis shows normalized exon reads (n = 10,383; read counts ≥ 10). Blue dots represent significantly changed genes in differential gene expression analysis (Wald test, adjusted p < 0.05). **D.** Venn diagram of mRNAs downregulated in cells expressing indicated Flag-SPT5 variants. **E.** Effects of expressing indicated Flag-SPT5 variants on representative genes induced by loss of CTR1 phosphorylation. **F.** Correlations of fold changes between different SPT5 variants in RNA-seq data. Pearson correlation R and p values are indicated. **G.** Violin plots of gene length for the indicated gene sets. Wilcoxon signed rank test, gene number (n) and p-value are indicated. **H.** Sashimiplot of significantly changed skipped exon (SE) events at the indicated gene locus. **I.** Venn diagram of indicated gene sets in cells expressing *muR1R2* mutant variant of Flag-SPT5.

**Figure S4.**
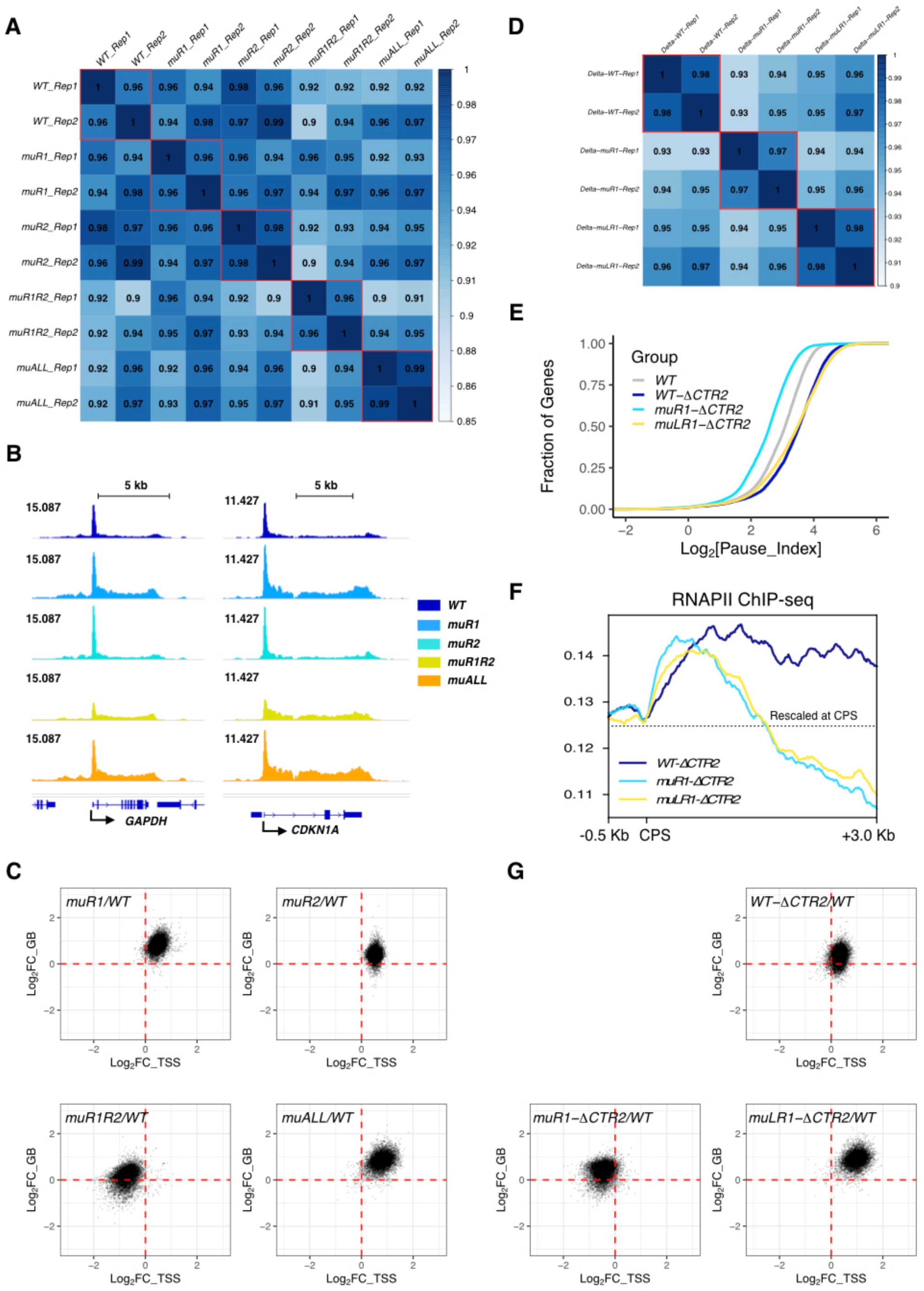
Control of pausing by tripartite SPT5 phosphorylation. Related to Figure 4. **A.** Reproducibility of RNAPII (RPB1) ChIP-seq data. Pearson correlations between replicates from *dTAG-SUPT5H* cells expressing indicated, full-length Flag-SPT5 variants. **B.** RNAPII ChIP-seq browser tracks of *GAPDH* and *CDKN1A* genes in cells expressing indicated, full-length Flag-SPT5 variants. **C.** Scatter plots of RNAPII occupancy fold change over transcription start site (TSS, *x*-axis) and gene body (GB, *y*-axis) in cells expressing indicated, full-length Flag-SPT5 variants (n = 8, 022). **D.** Pearson correlations between replicates of RNAPII ChIP-seq data from cells expressing indicated Flag-SPT5 variants with internal deletions of CTR2. **E.** ECDF plots of pausing index for genes with high RNAPII occupancy (n = 10,015) in cells expressing indicated Flag-SPT5 variants. Pausing index plot in cells expressing *WT* variant from Figure 4D is added for comparison. **F.** CPS-centered metagene plot of RNAPII ChIP-seq data from cells expressing indicated Flag-SPT5 variants with internal deletions of CTR2. Data were re-scaled at CPS. **G.** Scatter plots of RNAPII occupancy fold change over transcription start site (TSS, *x*-axis) and gene body (GB, *y*-axis) in cells expressing indicated Flag-SPT5 variants with internal deletions of CTR2 (n = 8, 019).

**Figure S5.**
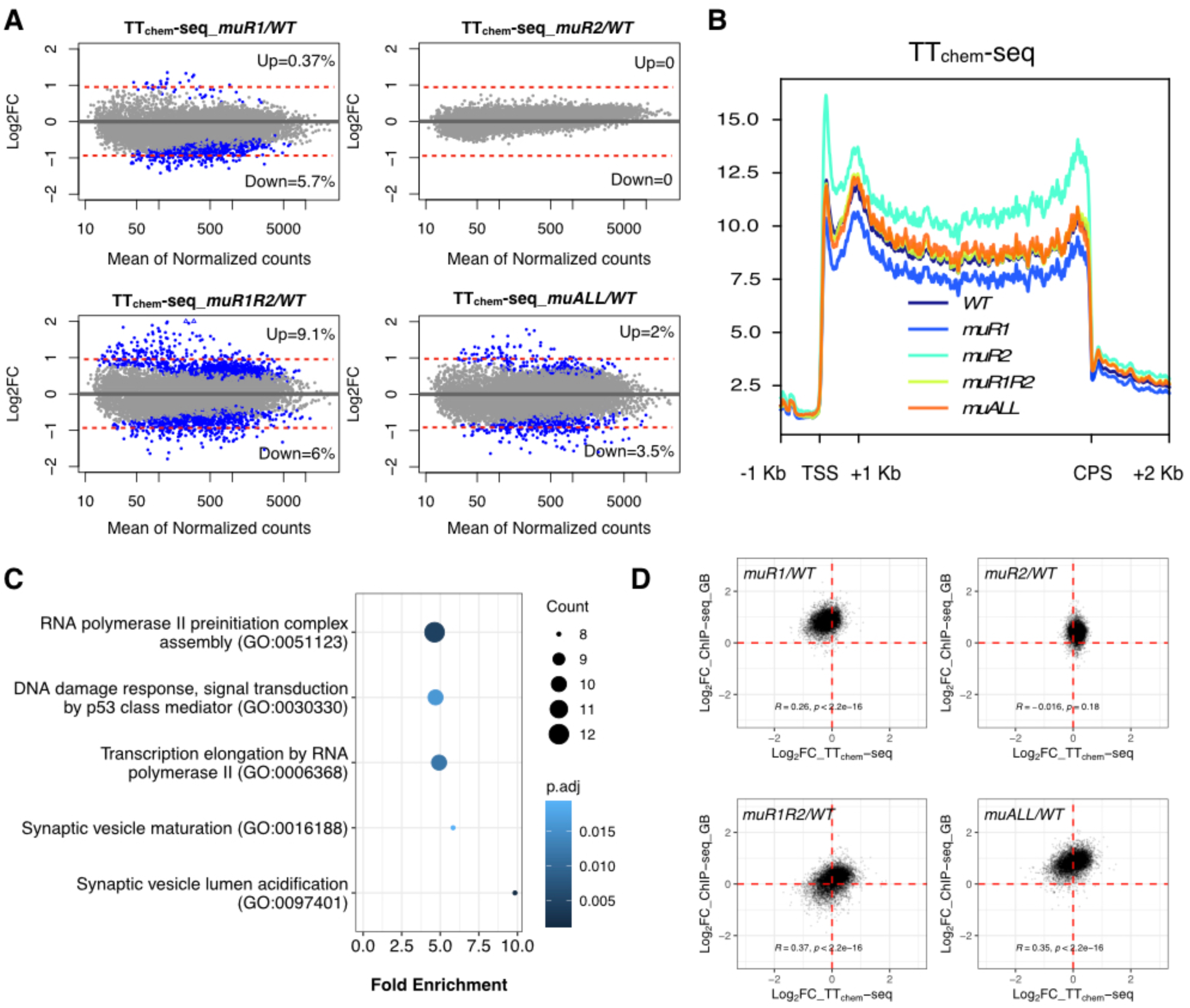
Combinatorial effects of CTR1 and CTR2 mutations on nascent transcription. Related to Figure 5. **A.** MA plots of TT_chem_-seq data. The *y*-axis shows log2 transformed fold changes and *x*-axis shows normalized exon reads (n = 11841; read counts>=10). Blue dots represent significantly changed genes in differential gene expression analysis (Wald test, adjusted p < 0.05). **B.** Merged metagene plots (sense strand) of TT_chem_-seq data from non-overlapping genes > 1 kb in length (n = 7,448). **C.** Gene Ontology (GO) term enrichment of nascent transcripts up-regulated in the double-CTR mutant. **D.** Scatter plots showing correlation between TT_chem_-seq (*x*-axis) and RNAPII ChIP-seq (*y*-axis) data in cells expressing indicated, full-length Flag-SPT5 variants. Pearson correlation R and p values are indicated.

**Figure S6.**
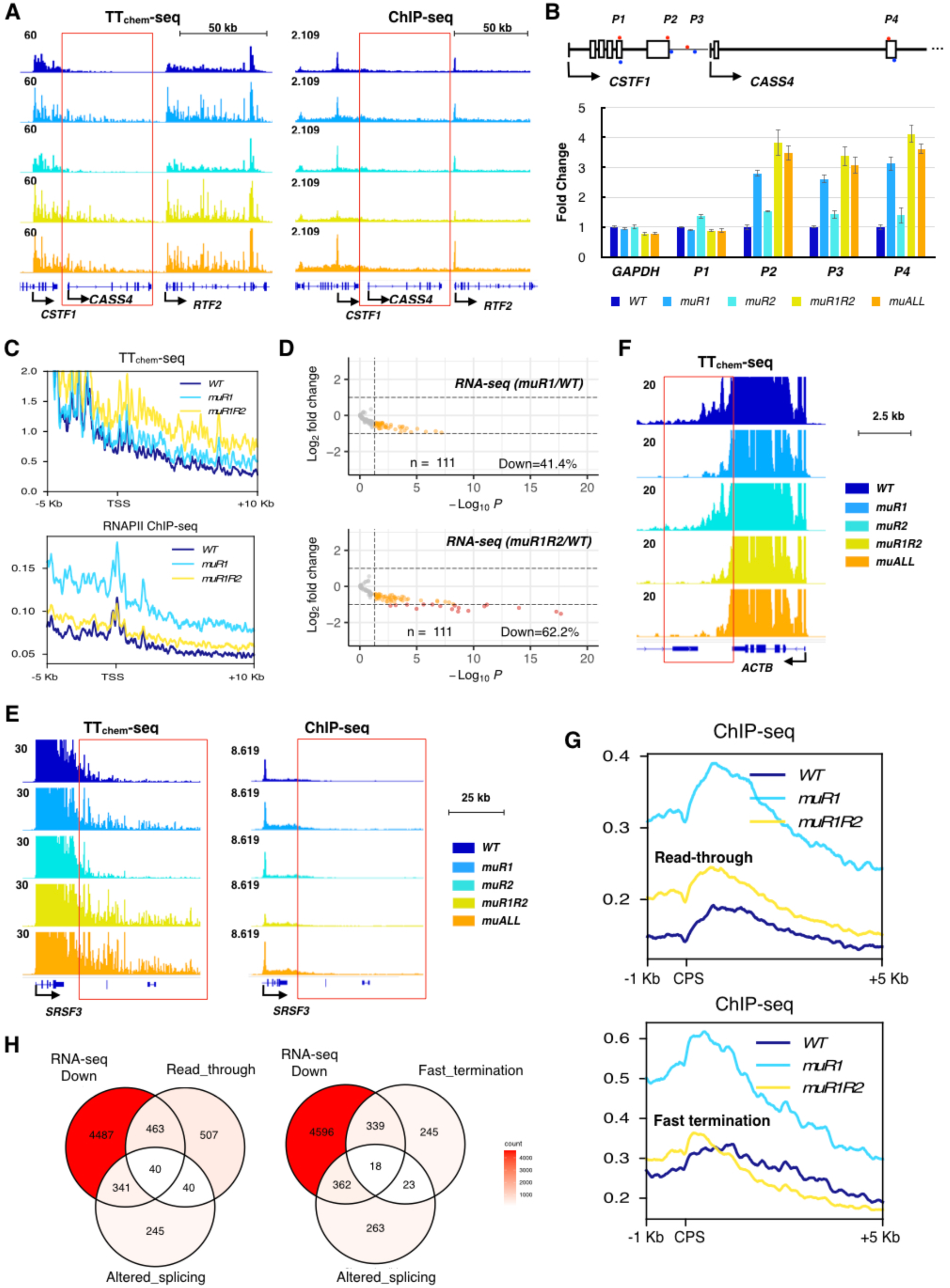
Combinatorial effects of CTR1 and CTR2 mutations on nascent transcription. Related to Figure 6. **A.** Browser tracks of TT_chem_-seq and RNAPII ChIP-seq data at *CASS4* locus. **B.** Verification of read-through transcription in cells expressing the double-CTR mutant variant by RT-qPCR with indicated primer pairs. Data are from three biological replicates (n = 3); error bars indicate mean ± SD. **C.** TSS-centered metagene plots (sense strand) of TT_chem_-seq and RNAPII ChIP-seq data from genes with low RNAPII occupancy that were upregulated by dual CTR mutation in TT_chem_-seq analysis (n = 159). **D.** Volcano plots of RNA-seq data for upstream genes with read-through transcription (n = 111). **E.** Browser tracks of TT_chem_-seq and RNAPII ChIP-seq data at *SRSF3* locus where transcriptional read-through occurs. Region downstream of CPS encompassing normal termination zone is demarcated by red box. **F.** Browser tracks of TT_chem_-seq data at *ACTB* locus, where faster termination occurs. Region downstream of CPS encompassing normal termination zone is demarcated by red box. **G.** CPS-centered metagene plots of RNAPII ChIP-seq data from indicated mutants at read-through (n = 1,307) and fast-termination genes in *muR1R2* (n = 866). **H.** Venn diagram of indicated gene sets in cells expressing *muR1R2* mutant of Flag-SPT5.

**Figure S7.**
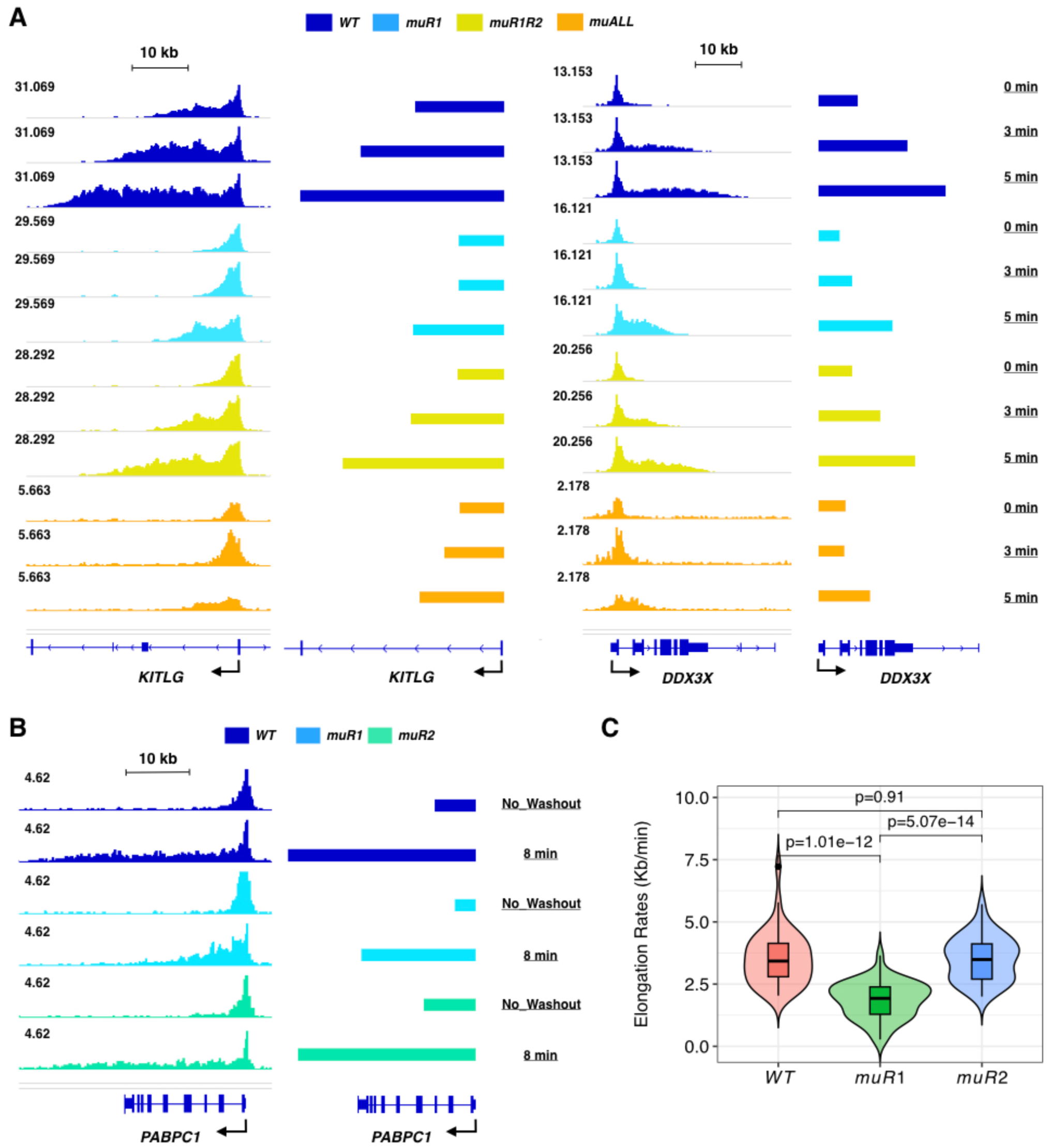
CTR1 and CTR2: accelerator and brake on RNAPII elongation. Related to Figure 7. **A.** Browser tracks from RNAPII (RPB1) CUT&RUN analysis on *KITLG* and *DDX3X* genes after DRB block and release for times indicated at right, in cells expressing indicated, full-length Flag-SPT5 variants: *WT*, *muR1* and *muR1R2*. Two biological replicates were analyzed (n = 2). **B.** Browser tracks of RNAPII CUT&RUN analysis on *PABPC1* gene after DRB block and release for times indicated at right, in cells expressing indicated, full-length Flag-SPT5 variants: *WT*, *muR1* and *muR2*. The data are from two biological replicates (n = 2) comparing only CTR1 and CTR2 single mutants to wild-type. **C.** Violin plot of elongation rates calculated for non-overlapping genes with high signal/noise ratio and > 30 kb in length (n = 41). Paired t-test, p values are as indicated.

Table S1. Excel file containing elongation rates determined by DRB washout RNAPII CUT&RUN, related to Figure 7.

Table S2. Excel file containing information on primers and synthesized DNA in this study.

Table S3. Excel file containing scale factors or size factors used for sequencing data.

## STAR METHODS

### KEY RESOURCES TABLE

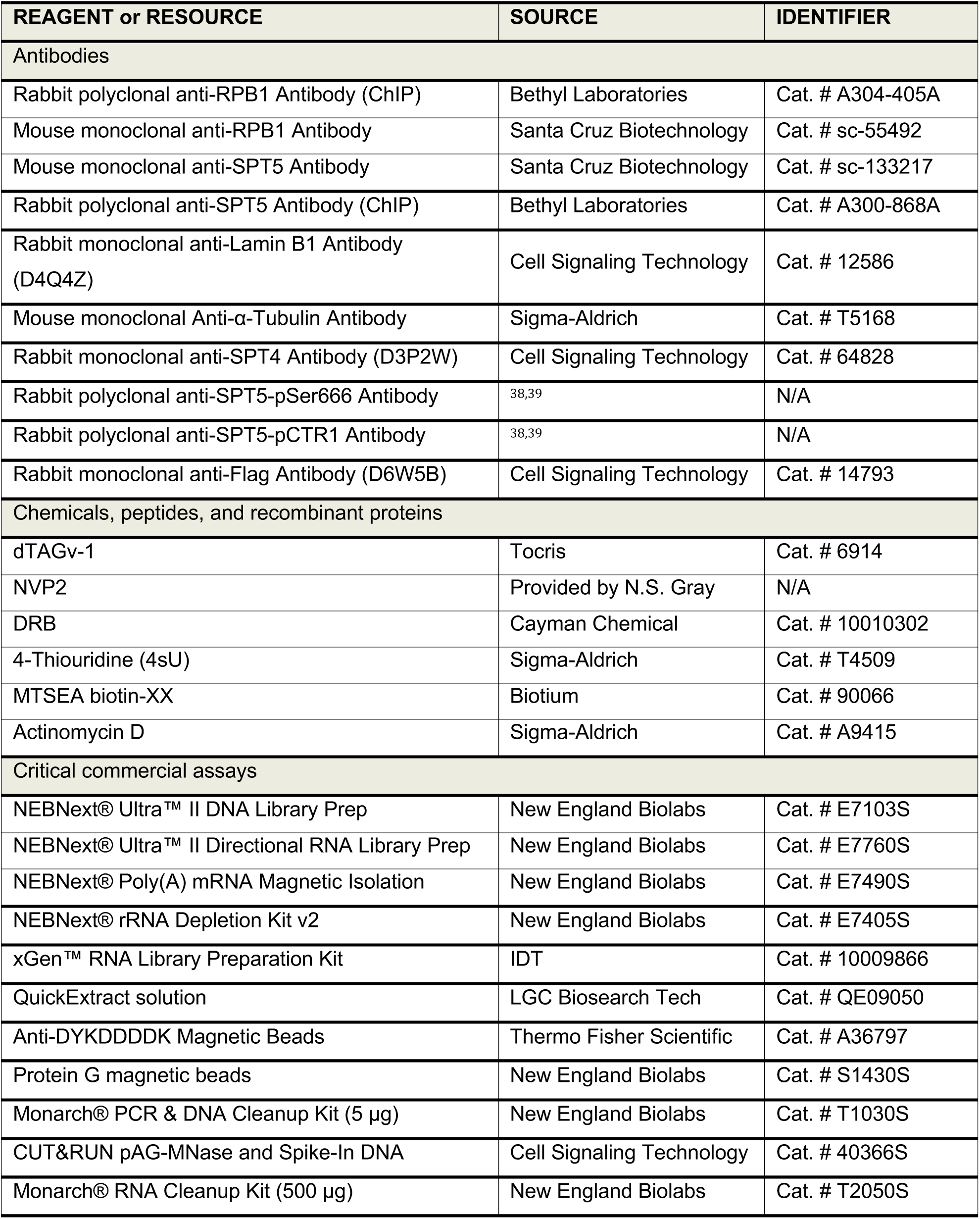
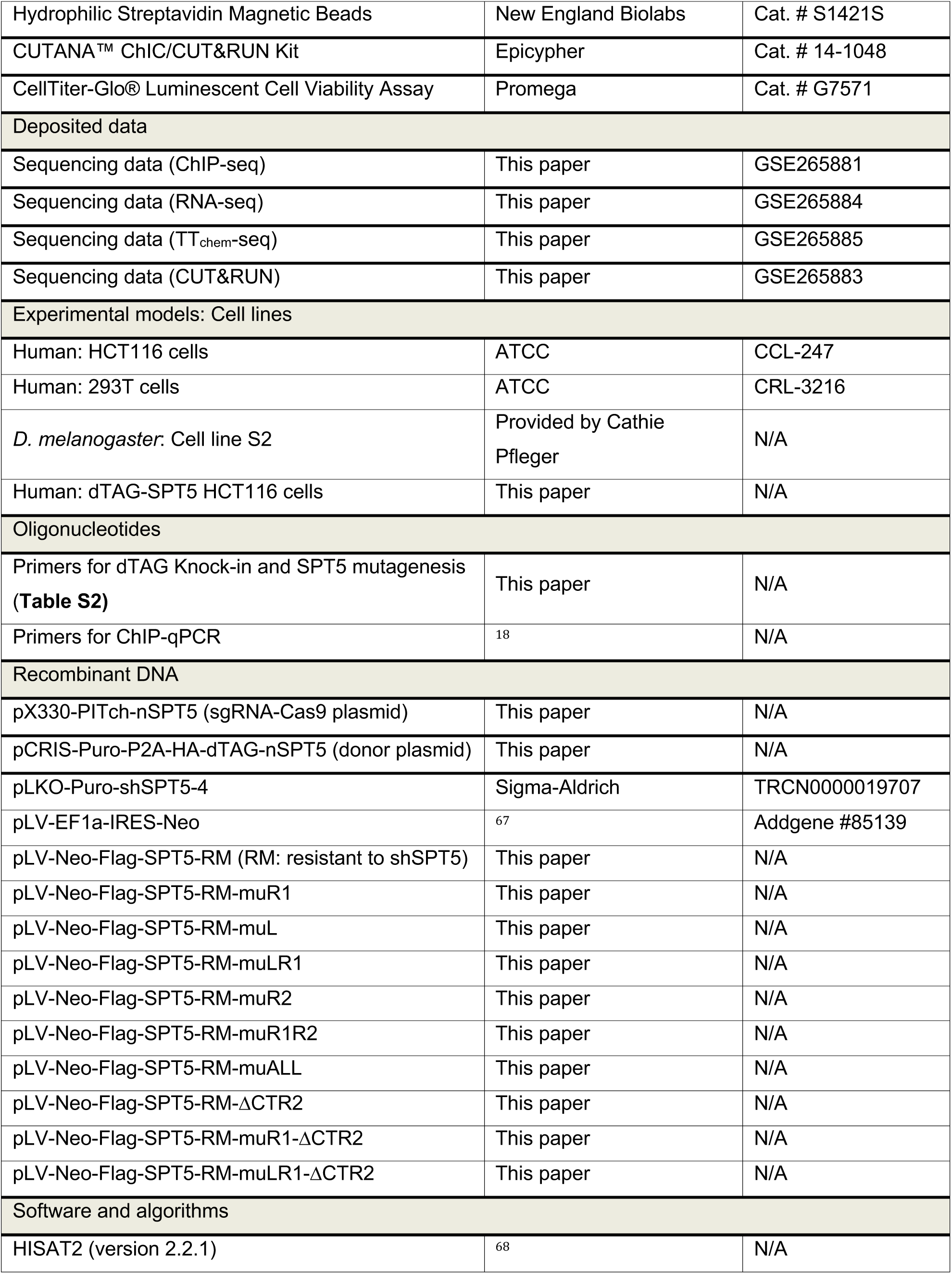
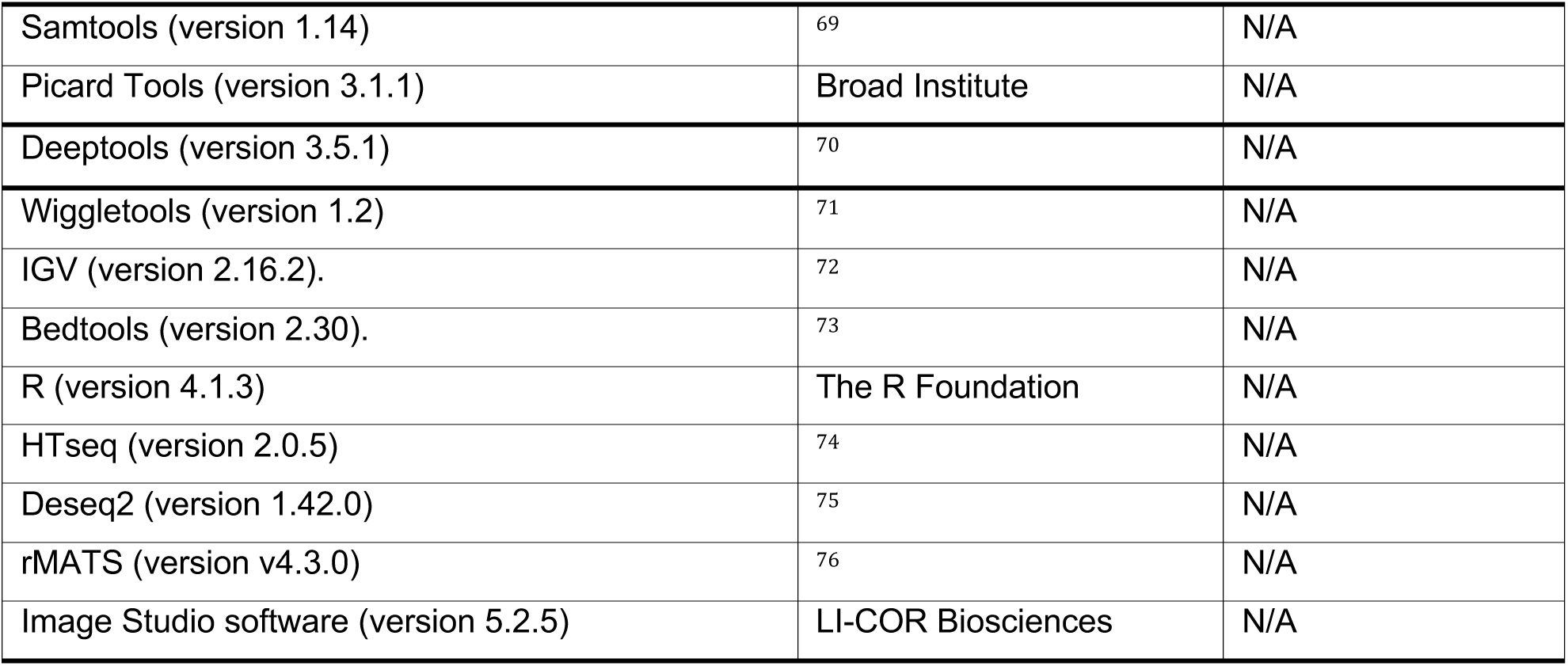

### RESOURCE AVAILABILITY

#### Lead contact

Requests for further information and reagents may be directed to and will be fulfilled by the lead contact, Robert P. Fisher

#### Materials availability

Unique reagents generated in this study are available from the lead contact.

#### Data and code availability

Sequencing data have been deposited at GEO (Accession numbers and tokens: GSE265881; GSE265883; GSE265884; GSE265885).

### EXPERIMENTAL MODEL AND SUBJECT DETAILS

#### Cell lines

HCT116 cells were cultured in McCoy’s 5A medium (Corning) supplemented with 10% Fetal Bovine Serum (FBS) (Gibco) and 1× Penicillin-Streptomycin (Corning) at 37°C with 5% CO_2_. 293T cells were cultured in DMEM medium (Corning) supplemented with 10% FBS and 1× Penicillin-Streptomycin at 37°C with 5% CO_2_. *Drosophila* S2 cells were cultured in Schneider’s medium (Gibco) supplemented with 10% inactivated FBS (Gibco) and 1× Penicillin-Streptomycin at 25°C.

### METHOD DETAILS

#### Construction of *dTAG-SUPT5H* cells

*dTAG-SUPT5H* HCT116 cells were generated by CRISPR-Cas9 genome editing with the PITCh system as previously described ^77,78^. Briefly, HCT116 cells grown in 6-well plates were co-transfected with 2 μg donor plasmid and 4 μg sgRNA-Cas9 plasmid with 20 μL lipofectamine2000 (Thermo Fisher Scientific). Cells were transferred to 10-cm dishes 1 day post-transfection to avoid confluence and selected with 0.8 μg/ml puromycin 4 days post-transfection. Cell clones resistant to puromycin were visible in 3 weeks. The genomic DNA of isolated clones was extracted with QuickExtract solution (LGC Biosearch Technologies) and tested by PCR. Clones with homozygous integration of the dTAG cassette at the *SUPT5H* locus were validated by Sanger sequencing and immunoblotting. To induce the degradation of SPT5 protein, 125 or 500 nM dTAGv-1 (Tocris) was added to the culture.

#### Lentivirus packaging and infection

To package lentivirus, HEK 293T cells grown in 10-cm dishes were co-transfected with 10 μg lenti-plasmid and 10 μg packaging plasmids (5 μg psPAX2 and 5 μg pMD2.G) with 60 μL 1 mg/mL polyethyleneimine. Lentivirus was collected 72 hr post-transfection and concentrated with 4x Lentivirus concentrator (40% PEG-8000, 1.2 M NaCl in PBS). Cells were infected with lentivirus in the presence of 10 μg/ml polybrene (Santa Cruz Biotechnology) and selected with 1 mg/ml G418 (Invivogen) or 1 μg/ml puromycin 72 hr post-infection.

#### Cell lysis and fractionation

Cells were fractionated as previously described ^79^ with minor modifications. Cells (8 × 10^6^ per 10-cm dish) were harvested by trypsinization and washed in PBS. Cells were resuspended without pipetting in 700 μL ice-cold CSK buffer (10 mM PIPES [pH 6.8], 100 mM NaCl, 300 mM sucrose, 3 mM MgCl_2_, 1 mM EGTA, 0.5% Triton X-100, 1 mM ATP) with protease and phosphatase inhibitors (1 mM PMSF, 5 μg/ml leupeptin, 2 μg/ml aprotinin, 50 mM NaF, 10 mM sodium β-glycerophosphate, 1 mM NaVO_3_) and incubated on ice for 10 min. The soluble fraction was collected by centrifugation at 1500 × g_av_ for 5 min at 4°C. The pellet was washed with CSK buffer once without pipetting and resuspended in 200 μL RIPA buffer (50 mM Tris-HCl [pH 7.4], 150 mM NaCl, 1% NP-40, 0.5% Na-deoxycholate, 0.1% SDS, 2 mM EDTA) with protease and phosphatase inhibitors. Lysates were sonicated in a Bioruptor (Diagenode) for 10 min with cycles of 30 s ON and 30 s OFF and clarified by centrifugation at 16,000 × g_av_ for 10 min at 4 °C. The supernatant was collected as the insoluble (chromatin) fraction. Alternatively, for experiments that did not require a soluble fraction, cells grown on 6- or 12-well plates were directly lysed in 1 mL ice-cold CSK buffer with protease and phosphatase inhibitors on ice for 15 min with shaking. The detached cells were transferred to tubes with wide-bore tips and processed as described above to obtain an insoluble fraction.

#### Immunoprecipitation and immunoblotting

For immunoprecipitation of Flag-tagged proteins, clarified lysates were incubated with 30 μL anti-Flag antibody-conjugated magnetic beads (Thermo Fisher Scientific) for 6 hr at 4 °C with end-to-end rotation. Beads were washed 4 times with ice-cold RIPA buffer. For immunoblot analysis, samples were separated by SDS-PAGE and transferred to 0.45-μm nitrocellulose membranes. The membranes were probed with primary antibodies and fluorescent second antibodies at 1:1000 and 1:5000 dilutions, respectively. Immunoblot signals were detected and quantified with Odyssey CLx Imaging System (LI-COR Biosciences) using Image Studio software (version 5.2.5).

#### Cell counting and viability assays

For cell counting, cells were seeded at different densities for measurement on different days. For cell viability assays, ∼2 × 10^3^ cells per well were seeded in 96-well plates and treated with 0.5 μM dTAGv-1 on the next day (Day 0). Four replicates were performed for each time point. Luminescent (white plate) or fluorescent (black plate) signals were recorded with a plate reader (BioTek) after adding CellTiter-Glo Reagent (Promega) or Resazurin working solution (10×, 0.01% in PBS).

#### ChIP-qPCR and ChIP-seq

ChIP was performed as previously described ^38^ with minor modifications. About 2 × 10^7^ HCT116 cells grown in 15-cm dishes were cross-linked with 1% formaldehyde for 10 min at room temperature. Cross-linking was quenched with 125 mM glycine for 5 min. Cells were collected with a cell lifter (Costar) in 2 ml ChIP nuclear fractionation buffer (50 mM HEPES [pH 7.5], 140 mM NaCl, 1 mM EDTA, 10% Glycerol, 1% NP-40, 0.25% Triton X-100) with protease and phosphatase inhibitors and incubated on ice for 10 min. Cell nuclei were collected by centrifugation at 600 x g_av_ for 10 min at 4 °C and washed with wash buffer (10 mM Tris-HCl [pH 8.0], 200 mM NaCl, 1 mM EDTA, 0.5 mM EGTA). The pellets were lysed in 1.8 mL RIPA buffer with protease and phosphatase inhibitors and sheared by sonication in a Bioruptor (Diagenode) at high power, for 2 x 20 min with cycles of 30 s ON and 30 s OFF. Lysates were clarified by centrifugation at 16,000 × g_av_ for 10 min at 4 °C. Cleared lysates (600 μL) were incubated with 4 μg antibody overnight at 4 °C with end-to-end rotation. Protein G magnetic beads (22 μL, New England Biolabs) were added to the tube on the second day and samples were incubated for another 4 hr at 4 °C with end-to-end rotation. The beads were washed 3 times with RIPA buffer, 3 times with IP wash buffer (100 mM Tris-HCl [pH 8.0], 500 mM LiCl, 1% NP-40, 1% sodium deoxycholate) and twice with TE buffer (10 mM Tris-HCl [pH 8.0], 1 mM EDTA). Cross-links were reversed by incubating beads in elution buffer (10 mM Tris-HCl [pH 8.0], 1 mM EDTA, 1% SDS, 200 mM NaCl, 0.8 U Proteinase K) at 55 °C in a block with heated lid for 16 hr. RNase A (1 μg) was added and samples were incubated at 37 °C for 30 min. DNA was purified with a DNA Cleanup Kit (5-μg scale, New England Biolabs). The purified DNA was used for qPCR and library preparation. *S. cerevisiae* genomic DNA (Cell Signaling Technology) was added to each sample for spike-in normalization and DNA libraries were prepared with NEBnext Ultra II DNA library kit (New England Biolabs). Two or more biological replicates were performed for each condition.

#### RNA-seq

Total RNA was extracted from cells with Trizol (Thermo Fisher Scientific). *Drosophila* RNA from S2 cells was added to samples (1:50) for spike-in normalization. NEBNext Poly(A) mRNA Magnetic Isolation Module (New England Biolabs) was used to select mRNA. RNA-seq library was prepared with xGen™ RNA Library Preparation Kit (IDT). Two or more biological replicates were performed for each condition.

#### TT_chem_-seq

TT_chem_-seq was performed as previously described ^59^. Briefly, ∼2 × 10^7^ cells were pulse-labeled with 0.5 mM 4-thiouridine (4sU, Sigma-Aldrich) for 15 min at 37 °C and lysed in 3 mL Trizol. Total RNA was extracted and 300 μg was used per experiment. *Drosophila* RNA from 4sU-labeled S2 cells was added to samples (1:50) for spike-in normalization. RNA was fragmented by adding 1 M NaOH to 0.2 M final concentration and incubating on ice for 20 min. Reactions were stopped by adding 1 M Tris buffer (pH 6.8) to 0.5 M final concentration. RNA samples were biotinylated with MTSEA biotin-XX (Biotium) and incubated in the dark for 30 min at room temperature. RNA was purified with RNA Cleanup Kit (500 μg-scale, New England Biolabs). Biotinylated RNA was enriched with hydrophilic streptavidin magnetic beads (New England Biolabs). Ribosomal RNA (rRNA) was removed with NEB rRNA removal module (New England Biolabs). Libraries were prepared with NEBNext Ultra II Directional RNA Library Prep Kit (New England Biolabs). Two or more biological replicates were performed for each condition.

#### CUT&RUN

Freshly prepared nuclei from ∼2 × 10^6^ cells were used per CUT&RUN experiment, performed according to published procedures ^80^. *E.coli* DNA (Epicypher) was added to each sample for spike-in normalization and DNA libraries were prepared with NEBnext Ultra II DNA library kit (New England Biolabs). For DRB-CUT&RUN, dTAG-SPT5 cells grown in 10-cm dishes were first treated with 125 nM dTAGv-1 for 4 hr, followed by addition of 100 μM DRB for a 3-hr incubation. DRB was washed out and cells were incubated in fresh medium with 125 nM dTAGv-1 at 37 °C for the indicated times. Cells were collected in medium containing 1 µg/ml of actinomycin D on ice, extracted and processed as decribed above for standard CUT&RUN. Two or more biological replicates were performed for each condition.

#### Next-generation sequencing and data processing

The concentration and size distribution of sequencing libraries were determined by Qubit (Invitrogen) and Tapestation or Bioanalyzer (Agilent). Libraries were sequenced on Illumina Hi-seq or NovaSeq (2×150) platforms. Reads were trimmed and filtered using TrimGalore (version 0.6.10) and aligned to the hg38 human genome (Ensembl) and spike-in genomes with HISAT2 (version 2.2.1). PCR duplicates were removed using Picard Tools (version 3.1.1) for ChIP-seq and TT_chem_-seq experiments. Alignment files were converted to Bam files with Samtools (version 1.14) for downstream analysis.

### QUANTIFICATION AND STATISTICAL ANALYSIS

#### ChIP-seq and CUT&RUN data analysis

Bigwig files were generated in 10-base pair (bp) bins with the BamCoverage function in Deeptools (version 3.5.1). Reads were normalized using the CPM method and scaled based on the spike-in/intergenic region signal (**Table S3**). The bigwig files of replicates in the experiment were averaged using wiggletools (version 1.2). Bigwig files were used to generate gene browser tracks in IGV (version 2.16.2). Data matrices were generated with the computeMatrix function in Deeptools. The matrix files were used to produce the metagene plots and heatmaps. Non-overlapping genes > 1 kb were selected with the intersect function in bedtools (version 2.30). The intergenic region was selected with the complement function in bedtools (gene-adjacent regions 10 kb upstream of TSS and 10 kb downstream of CPS were not included). To analyze the promoter-proximal region (TSS to +300 bp) and gene body (+300 bp to CPS) separately, the mean signals in both regions of each gene were separately calculated with the multiBigwigSummary function in Deeptools. Empirical cumulative distribution function (ECDF) plots of RNAPII pausing index and fold-change scatterplots were generated in R (version 4.1.3). The Pearson correlations between samples were calculated with the plotCorrelation function in Deeptools and re-generated in R.

#### RNA-seq data analysis

Reads mapped to exons and introns were counted with HTseq (version 2.0.5). Differential gene expression analysis (exon reads) was performed with Deseq2 (version 1.42) in R. The sizeFactors in DEseq2 were adjusted based on the spike-in alignment ratio. Deseq2 uses the Wald test to identify differentially expressed genes by default. The threshold of adjusted p value was set at 0.05. Alternative splicing analysis was performed with rMATS (version v4.3.0).

#### TT_chem_-seq data analysis

Reads mapped to introns, exons, and genes were counted with HTseq (version 2.0.5). Differential gene expression analysis (intron reads) was performed with Deseq2 (version 1.42.0) in R. The sizeFactors in DEseq were adjusted based on the spike-in alignment ratio. BedGraph files were generated with the genomecov function in bedtools (version 2.30). Reads were normalized and scaled based on the sizeFactors used in DEseq2.

BedGraph files were converted into bigwig files for later analysis. The bigwig files of replicates were averaged using wiggletools (version 1.2). Data matrices were generated with the computeMatrix function in Deeptools (version 3.5.1). The matrix files were used to produce metagene plots. The elongation index was calculated by computing the ratio between TT_chem_-seq signal and ChIP-seq signals with wiggletools. Metagene plots of elongation index were generated as described above. To analyze the signal downstream of the CPS (5-kb window), the mean signal in the window for each gene was calculated using wiggletools.

#### Calculation of elongation rates

To calculate elongation rates, the bigwig files from DRB CUT&RUN experiments were divided into 100-bp bins through the bin function of wiggletools. Non-overlapping, highly expressed genes longer than 30 kb were selected by using the intersect function in bedtools. Local background was set through gamma distribution modeling (maximum likelihood method) and quantile method for each gene by using the eqgamma function in the EnvStats package of R. Regions with signal above the local background were extracted and merged if the distance between features was within 500 bp (5 bins) by using the merge function in bedtools. The RNAPII wave regions were selected in R by matching regions that start at the TSS. Genes with typical RNAPII wave profiles (wavefront edge moving forward with time) were selected for elongation rate calculation. The RNAPII wave bed files were examined in the IGV gene browser to ensure the accuracy of the RNAPII wavefront edge. The early, late, and averaged elongation rates were calculated by using the distances between the wavefront edges of 0-3 min, 3-5 min and 0-5 min time intervals, respectively. Paired t-test was used to compare the elongation rates between samples. Null hypothesis: no significant difference in mean elongation rates between samples.

